# Repairing a deleterious domestication variant in a floral regulator gene of tomato by base editing

**DOI:** 10.1101/2024.01.29.577624

**Authors:** Anna N. Glaus, Marion Brechet, Gwen Swinnen, Ludivine Lebeigle, Justyna Iwaszkiewicz, Giovanna Ambrosini, Irene Julca, Jing Zhang, Robyn Roberts, Christian Iseli, Nicolas Guex, José Jiménez-Gómez, Natasha Glover, Gregory B. Martin, Susan Strickler, Sebastian Soyk

## Abstract

Crop genomes accumulated deleterious mutations, a symptom known as the cost of domestication. Precision genome editing has been proposed to eliminate such potentially harmful mutations, however, experimental demonstration is lacking. Here, we identified a deleterious mutation in the tomato transcription factor *SUPPRESSOR OF SP2* (*SSP2*), which became prevalent in the domesticated germplasm and diminished DNA-binding to genome-wide targets. We found that *SSP2* acts partially redundant with its paralog *SSP* to regulate shoot and inflorescence architecture. However, redundancy was compromised during tomato domestication and completely lost in the closely-related species *Physalis grisea*, in which a single ortholog regulates shoot branching. We applied base editing to directly repair the deleterious mutation in cultivated tomato and obtained plants with compact growth that provide an early fruit yield. Our work shows how deleterious variants sensitized modern genotypes for phenotypic tuning and illustrates how repairing deleterious mutations with genome editing may allow predictable crop improvement.

## MAIN

Deleterious mutations lead to the alteration or loss of gene activity. Crop domestication has been accompanied by an accumulation of potentially deleterious mutations^1,2^, a phenomenon described as the genetic cost of domestication^3^. Such potentially harmful variants likely influence many important agricultural traits^4^. For example, harmful recessive alleles can have detrimental effects that are exposed in homozygous progeny during inbreeding^5^. Deleterious mutations are often considered to mainly negatively affect fitness of natural populations but recently, a more nuanced view has been proposed that considers their adaptive value^6,7^. Deleterious, loss-of-function mutations may confer an evolutionary advantage during rapid shifts in environmental conditions and the selective pressures thereof^7^. Crop domestication created novel environments under which many traits that were beneficial in the wild likely became neutral or even detrimental. Illustrative examples include loss of photoperiodic flowering and seed shattering. These observations support the “less-is-more” idea, which proposes that selection may favor a less-than-complete repertoire of functional genes^7^. Nonetheless, eliminating deleterious variants from domesticated germplasm has been proposed as a major goal in future crop breeding to avert potential harmful effects^4,8^. However, correcting genetic variants by recombination during cross-breeding can be complicated by genetic linkage with beneficial alleles or near fixation in domesticated populations. Recent advances in precision genome editing promise to facilitate the repair of deleterious variants^9^. However, to our knowledge, an experimental demonstration of precision genome editing for the repair of deleterious variants in domesticated germplasm has been lacking.

A recurrent target of selection during crop domestication and breeding are alterations in flowering time^9^. Changes in flowering time allowed the adaptation of crops to novel environments and growing seasons different from their wild ancestors’ origin. The floral transition also influences plant architecture by balancing vegetative and reproductive growth^10^. At the molecular level, flowering occurs when the universal flowering hormone, florigen, reaches a critical level that triggers stem cells in the shoot meristems to switch from vegetative to reproductive growth. In the model crop tomato (*Solanum lycopersicum*), florigen is encoded by *SINGLE FLOWER TRUSS* (*SFT*), a homolog of Arabidopsis *FLOWERING LOCUS T* (*FT*) and member of the *CENTRORADIALIS, TERMINATING FLOWER1, SELF-PRUNING* (*CETS*) gene family^11^. While *SFT* promotes the floral transition, *SELF PRUNING* (*SP*) acts as antiflorigen and opposes the activity of florigen to repress flowering^12^. Evidence from rice and Arabidopsis suggests that florigen protein competes with antiflorigen protein for Group-A basic region/leucine zipper (bZIP) transcription factors to form the Florigen Activation Complex (FAC)^12–14^. In tomato, the bZIP transcription factor SUPPRESSOR OF SP (SSP) is a functional FAC component and *ssp* mutations have been used to fine-tune plant architecture for optimized fruit productivity^15^. In other crops, mutations in central florigen pathway components have been also selected to change flowering time and shoot architecture^9^. Yet, how deleterious mutations affected key components of the florigen pathway during crop domestication has not been systematically studied.

## RESULTS

### Prediction of deleterious variants in the florigen pathway

To determine the mutational load in domesticated tomato, we generated a chromosome-scale genome assembly for the closely-related wild tomato species *S. pimpinellifolium* (accession LA1589) (see **Online Methods**). We used this wild tomato genome as a reference to identify nonsynonymous mutations across a collection of 82 genomes along the domestication history of tomato, including 27 wild tomato species (*S. pimpinellifolium*), 23 landrace (*S. lyc. var. cerasiforme*), and 32 domesticated (*S. lycopersicum var. lycopersicum*) genomes (**Fig. 1a**, **Supplementary Table 1)**^16,17^. We predicted deleterious variants by amino acid conservation modelling and identified 39,132 (23.1 %) nonsynonymous variants with a putative deleterious effect (SIFT-score < 0.05) (**Supplementary Fig. 1a, b, Supplementary Table 2**)^17^. This analysis indicated that wild species, landrace, and domesticated tomato genomes contain on average 5,114, 7,131, and 8,233 homozygous deleterious variants, respectively (**Supplementary Fig. 1c**). Next, we focused on core components of the FAC^14^ and searched for deleterious variants in *CETS* and Group-A bZIP genes (**Supplementary Fig. 2**)^18^. Among all 12 annotated tomato *CETS* genes, we identified three genes with predicted deleterious variants (**Fig. 1b**). Besides a predicted deleterious variant with low confidence in an uncharacterized *TERMINATING FLOWER1* (*TFL1*)-like gene, we detected high-confidence predicted deleterious variants in an uncharacterized *MOTHER OF FT* (*MFT*)-like gene (Solyc03g119100) and in the known flowering repressor *SELF-PRUNING 5G* (*SP5G*; Solyc05g053850). The deleterious variants in *MFT-like* and *SP5G* were absent from wild tomato genomes but detected in 6 (26%) and 13 (57%) landrace, and 30 (94%) and 32 (100%) domesticated genomes, respectively (**Fig. 1c**, **Supplementary Table 3**). We also detected the *sp-classic* breeding mutation (P76L) that was predicted to not be deleterious but tolerated, which supports a hypomorphic nature of the mutation^19^. Among all 13 annotated tomato Group-A bZIP genes, we identified two uncharacterized abscisic acid responsive element binding factor (*ABF*)-like genes and an SSP-like gene (Solyc02g061990) with high-confidence predicted deleterious mutations (**Fig. 1d and Supplementary Fig. 2**). The predicted deleterious variant in Solyc02g061990 was the most frequent and absent from wild tomato genomes but detected in 5 landrace (22%) and 31 domesticated (97%) genomes (**Fig. 1e**). We concluded from these analyses that several central florigen pathway components have acquired potentially deleterious mutations during tomato domestication.

**Figure 1:**
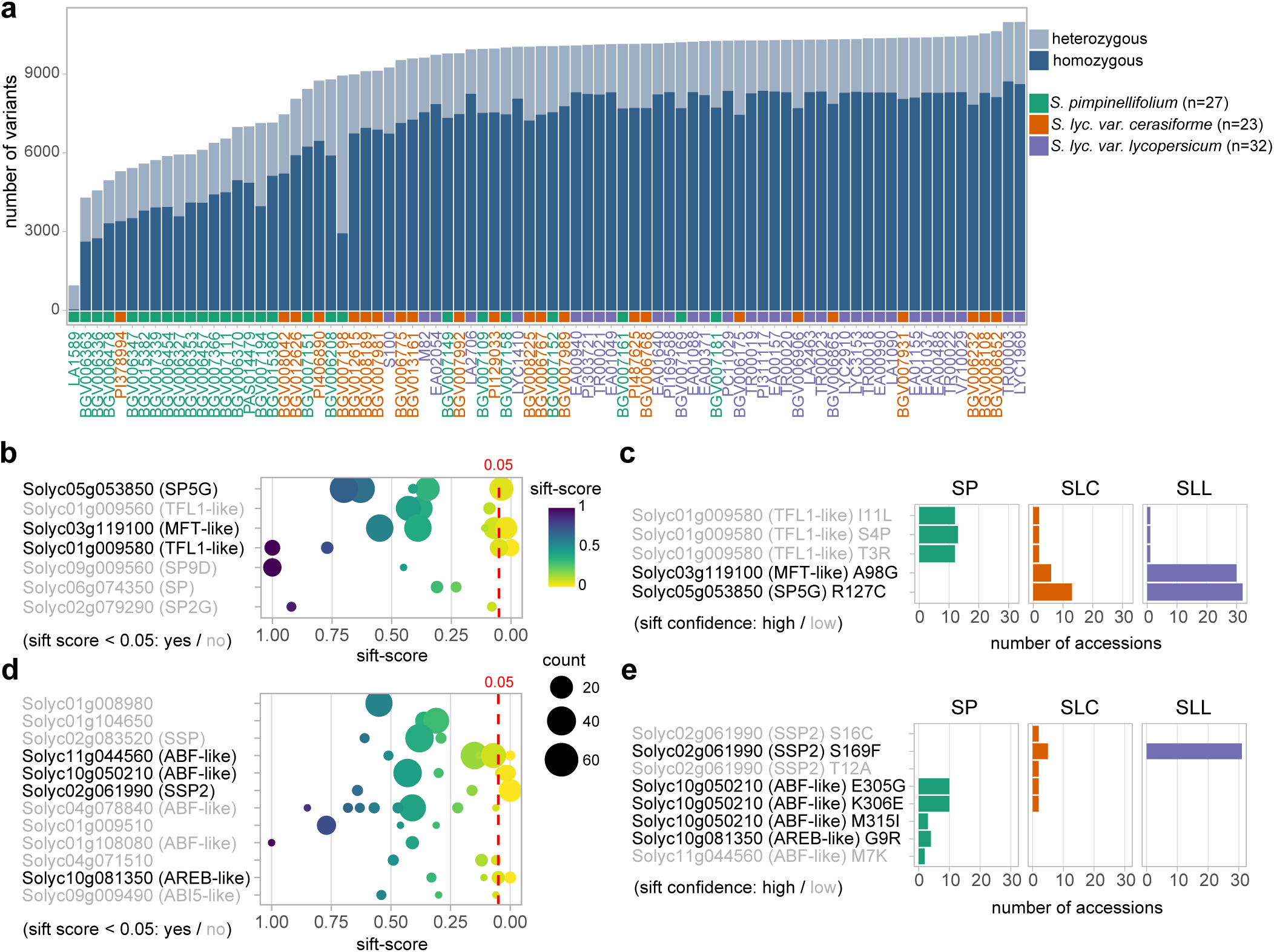
Predicting the load of deleterious variants along the domestication history of tomato. **a**, Number of predicted deleterious mutations in a panel of 82 tomato genomes, including wild species (*S. pimpinellifolium*, green), landraces (*S. lycopersicum var. cerasiforme*, orange), and cultivars (*S. lycopersicum*, purple). **b**, Prediction of deleterious variants across all annotated *CETS* genes. The dashed red line indicates the threshold for deleterious prediction (SIFT-score<0.05). Dot size scales with the number of genomes that carry the variant. Black and grey font indicates genes with predicted deleterious and tolerated mutations, respectively. **c**, Distribution of predicted deleterious *CETS* gene variants across wild (*S. pimpinellifolium*; SP), landrace (*S. lyc var. cerasiforme*; SLC) and domesticated (*S. lycopersicum*, SLL) genomes. Black and grey font indicate high and low prediction confidence, respectively. **d,e**, Prediction of deleterious variants in all annotated Group-A bZIP genes (d) and their distribution across different genome categories (e) as in (b, c).

### A missense mutation in *SSP2* accumulated during domestication

A phylogenetic analysis comparing group-A bZIP proteins of tomato and Arabidopsis showed that Solyc02g061990 is most closely related to *SSP*, thus we named the gene *SSP2* (**Fig. 2a** and **Supplementary Fig. 2**). SSP and SSP2 form a sister clade to the Arabidopsis proteins FD and FD PARALOG (FDP) ^20^, with SSP and FD being the more ancient genes. In Arabidopsis, FD and FDP are involved in flowering control and phytohormone responses^21,22^. Expression data from different tomato plant tissues showed that *SSP* and *SSP2* had similar expression patterns, suggesting functional redundancy, most notably in secondary (sympodial) shoot meristems (**Fig. 2b**)^23,24^. The putative deleterious variant in SSP2 causes a serine-to-phenylalanine (S169 to F169) exchange at a conserved residue in the DNA-binding domain (**Fig. 2c**). We analyzed the distribution of the ancestral (*SSP2^S169^*) and domesticated (*SSP2^F169^*) variants across 768 re-sequenced tomato accessions and found that the domesticated allele was absent from wild tomato species. The putative deleterious variant first arose in tomato landraces (*S. lycopersicum var. cerasiforme)*, was enriched in domesticated genotypes, and nearly fixed in modern fresh-market and processing types (**Fig. 2d)**. To genetically test if the putative deleterious variant has an effect on the floral transition, we introgressed the ancestral *SSP2^S169^* allele into a processing tomato type (cv. M82). We found that near-isogenic lines (NILs) harboring *SSP2^S169^* flowered earlier on sympodial shoots and developed shoots that grew more compact compared to the wild-type (WT) controls (**Supplementary Fig. 3a-f**). In addition, we introduced *SSP2^S169^* into the hypomorphic *ssp^2129^* mutant^15^ to test whether *SSP2^S169^* acts redundantly with its paralog *SSP*. We found that *SSP2^S169^* suppressed late-flowering and indeterminate growth of *ssp^2129^* mutants (**Supplementary Fig. 3g, h**), suggesting that the ancestral *SSP2^S169^* allele can compensate for reduced *SSP* activity.

**Figure 2:**
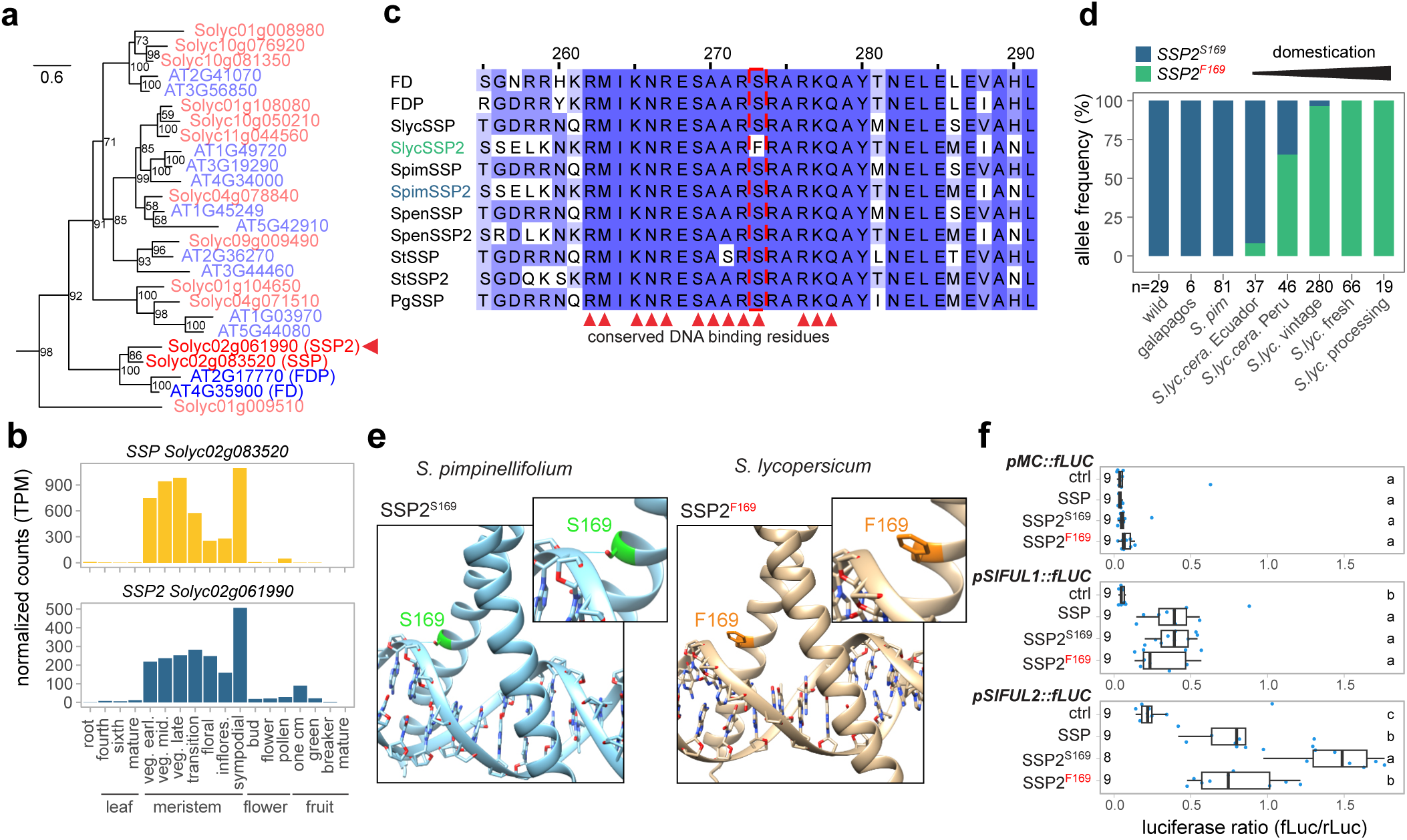
A deleterious mutation in *SSP2* reduces its transcription factor activity. **a**, Maximum-likelihood tree of A-group bZIP proteins in tomato (red font) and Arabidopsis (blue font). Red arrowhead marks SSP2. Numbers represent bootstrap values from 1,000 replicates and scale bar indicates the average number of substitutions per site. **b**, Normalized gene expression (TPM) for *SSP* and *SSP2* in different tissues and developmental stages (veg. earl./mid./late, stand for early, middle and late vegetative meristem stage). **c**, Partial alignment of SSP-like bZIP proteins from Arabidopsis, domesticated tomato (*S. lycopersicum*; *Slyc*), close wild tomato relative (*S. pimpinellifolium*; *Spim*), distant wild tomato relative (*S. pennellii*; *Spen*), potato (*S. tuberosum*; *St*), and *Physalis grisea* (*Pg*). Red arrowheads mark conserved DNA-binding residues. **d**, Distribution of ancestral (*SSP2^S169^*) and derived (*SSP2^F169^*) *SSP2* alleles in distant wild tomato relatives, wild relatives (*S. galapagense* / *S. cheesmaniae*), wild progenitor species (*S. pimpinellifolium*), landraces (*S. lyc var. cerasiforme*), and cultivars (*S. lycopersicum*). n=number of accessions. **e**, Predicted structures of ancestral SSP2^S169^ and derived SSP2^F169^ proteins on target DNA determined by homology modelling. Insets show a magnified view of the serine/phenylalanine residue at position 169. **f**, Reporter assays in tobacco leaves using SSP, SSP2^F169^, and SSP2^S169^ as effectors and firefly Luciferase (fLuc) driven by upstream sequences of *MC* (*pMC::fLUC*), *SlFUL1* (*pSlFUL1::fLUC*), and *SlFUL2* (*pSlFUL1::fLUC*) as reporter. Numbers indicate biological replicates (transformed leaves). Ctrl indicates the no effector control (35S::YFP). Letters represent the results from pairwise comparisons of means using one-way ANOVA and post-hoc Tukey’s HSD test with 95% confidence level. For box plots in (f), the bottom and top of boxes represent the first and third quartile, respectively, the middle line the median and the whiskers the maximum and minimum values.

### Domesticated SSP2 is compromised as a transcription factor

We hypothesized that the loss of the conserved serine residue affects the ability of SSP2 to bind DNA during the regulation of target genes. We modelled the structure of the ancestral (SSP2^S169^) and domesticated (SSP2^F169^) proteins in a homology-based modelling approach^25,26^. The model predicted that the conserved serine (S169) most likely forms hydrogen bonds with the phosphate backbone of the DNA target sequence whereas a phenylalanine at this position (F169) might increase the distance between the protein and target DNA due to its larger side-chain and hydrophobicity (**Fig. 2e**). To test whether the amino acid exchange affects the transcription factor function of SSP2, we co-expressed SSP2^F169^, SSP2^S169^ and SSP with SFT in tobacco leaves to quantify their transactivation activity on the upstream regions of *MACROCALYX* (*MC*; Solyc05g056620), *S. lycopersicum FRUITFULL1* (*SlFUL1*, Solyc06g069430), and *SlFUL2* (Solyc03g114830). These genes are homologous to Arabidopsis *APETALA1* and *FRUITFULL*, which have been shown to be activated by FD during the floral transition^13^. None of the effector constructs activated the *MC* reporter, which may result from a non-direct relationship between tomato *MC* and Arabidopsis *AP1*. In addition, we observed only low or nonsignificant transactivation of the *SlFUL1* reporter. However, the *SlFUL2* reporter was significantly activated by SSP and both SSP2 variants, and transactivation was significantly higher by the ancestral SSP2^S169^ compared to the SSP2^F169^ variant (**Fig. 2f, Supplementary Fig. 3i**). Together with the structural and genetic analyses, the results from this heterologous expression system suggested that the deleterious variant in *SSP2* disrupts the DNA-binding domain of domesticated SSP2^F169^ and compromises its transcription factor function.

To determine whether the deleterious SSP2^F169^ variant affects binding at genome-wide targets, we performed DNA affinity purification sequencing (DAP-seq) with SSP, ancestral SSP2^S169^ and domesticated SSP2^F169^ as bait proteins^27^. We identified 14,091 DAP-seq peaks that were significantly enriched (log_2_FC ≥ 3, FDR ≤ 0.01) compared to the input controls (**Fig. 3a and Supplementary Table 4**). The majority (7,388) of peaks were shared between SSP and the ancestral SSP2^S169^ but only 1,285 peaks were also bound by domesticated SSP2^F169^. We analyzed the genome-wide distribution of peaks for all three transcription factors and found more than 50% of peaks within proximal regulatory regions (**Fig. 3b**). *De-novo* motif enrichment analysis identified a G-box motif (CACGTG) with a subtle variation for SSP2^F169^ outside the core-motif (**Fig. 3c**). Next, we analyzed genes with proximal peaks (≤ 3 Kbp upstream and ≤ 2 Kbp downstream) and identified 7,688 and 5,010 putative target genes for SSP and SSP^S169^, of which the majority (4,699 genes) were bound by both proteins (**Fig. 3d** and **Supplementary Table 5**). In contrast, domesticated SSP2^F169^ bound only 1,175 and 1,136 of SSP and SSP^S169^ targets, respectively, and 1,616 genes in total. The low number of SSP2^F169^ targets and shared targets with SSP and SSP2^S169^ suggested that the ability of SSP2^F169^ to bind its genome-wide targets is compromised. To support this finding, we quantified binding intensity at target regions based on normalized read coverage. While SSP and SSP2^S169^ displayed similar binding intensities, SSP2^F169^ binding was strongly reduced (**Fig. 3e, f** and **Supplementary Fig. 4c, e**). Next, we conducted a differential binding analysis between SSP2^S169^ and SSP2^F169^ (see Methods). Of the 9,199 SSP2 peaks, we identified 5,063 differentially bound by SSP2^S169^ and SSP2^F169^ (log_2_FC ≥ 3, FDR ≤ 0.01) (**Supplementary Fig. 5a**). The majority (4,637) of differentially-bound peaks were private to SSP2^S169^ while only 426 private peaks were identified for SSP2^F169^ (**Supplementary Fig. 5b**). Private peaks for SSP2^S169^ were enriched for a G-box while a more relaxed TA-rich motif was enriched in SSP2^F169^ private peaks (**Supplementary Fig. 5c**). In addition, SSP2^F169^ private peaks tended to have a lower coverage, suggesting compromised binding ability also at private peaks (**Supplementary Fig. 5d,e**). In support of this, we did not detect any enriched gene ontology (GO) terms for the 230 genes near SSP2^F169^ private peaks while the 3,046 genes near SSP2^S169^ private peaks were enriched for genes related to circadian rhythm and rhythmic processes (**Supplementary Fig. 5f,g**). Differential binding of SSP2^F169^ and SSP2^S169^ was also obvious at the level of individual genes. For example, we found that the upstream regions of the two tomato homologs of *GIGANTEA* (*GI*), which regulates photoperiodic flowering in Arabidopsis^28^, were bound by SSP and SSP2^S169^ but not by the domesticated SSP2^F169^ variant (**Fig. 3g, h**). Together, our genome-wide binding data demonstrates that SSP and the ancestral SSP2^S169^ variant bind a set of largely shared targets while domesticated SSP2^F169^ is compromised in its ability to bind the targets of the ancestral protein.

**Figure 3:**
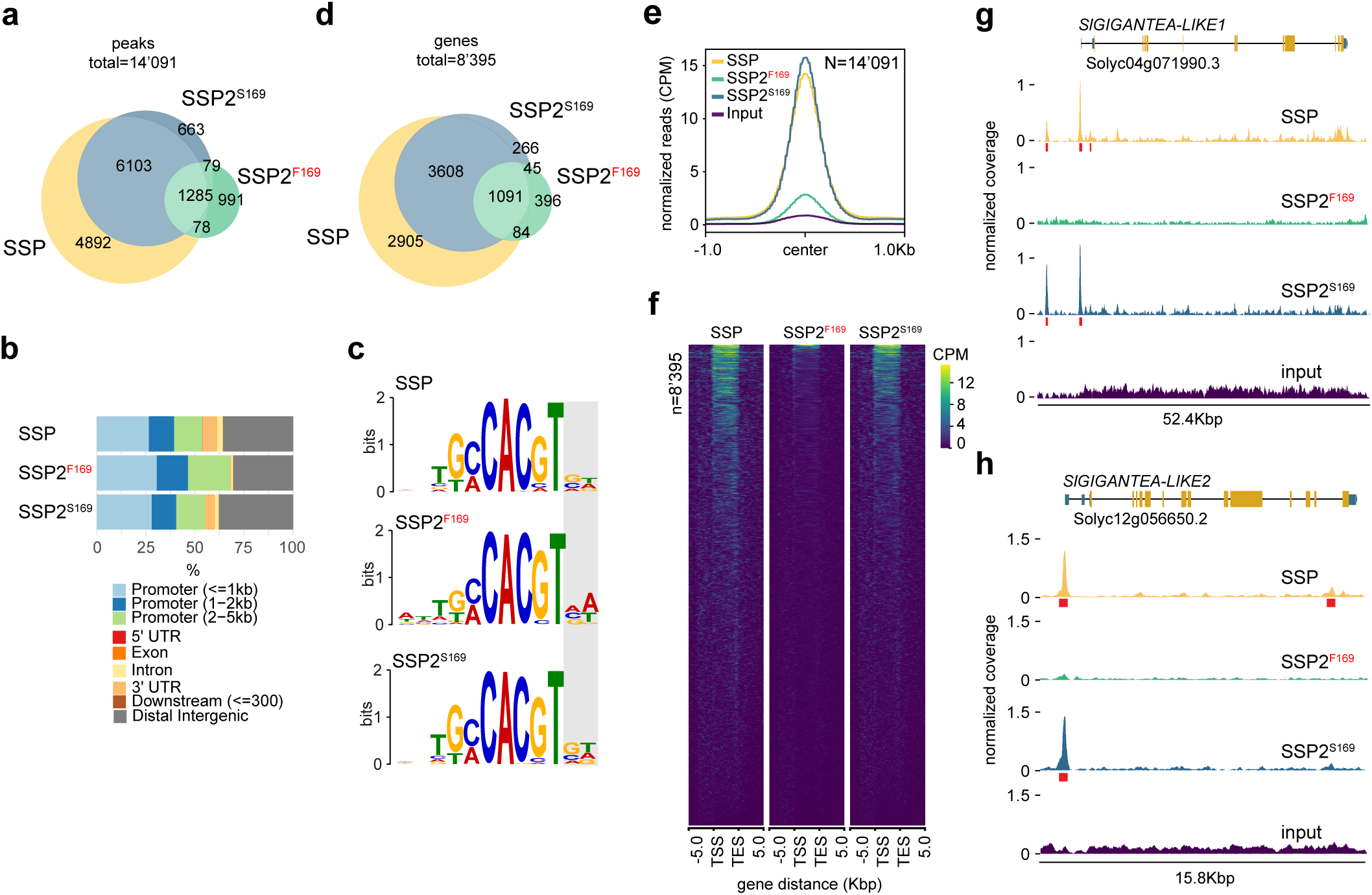
Domesticated SSP2^F169^ shows reduced binding at genome-wide target loci. **a**, Overlap of significant (log_2_FC ≥ 3, FDR ≤ 0.01) SSP, SSP2^F169^, and SSP2^S169^ DAP-seq peaks (n=14’091). **b**, Distribution of significant SSP, SSP2^F169^, and SSP2^S169^ DAP-seq peaks across gene features. **c**, Most-significant motifs identified by *de-novo* motif enrichment analysis of SSP, SSP2^F169^, and SSP2^S169^ DAP-seq peak regions. Grey box delimits region with motif variation outside the core-motif. **d**, Overlap of genes with significant DAP-seq peaks ≤ 3 Kbp upstream and ≤ 2 Kbp downstream of the transcriptional start site (n=8’395). **e**, Profiles of normalized read coverage at significant SSP, SSP2^F169^, and SSP2^S169^ peaks. **f**, Comparison of SSP, SSP2^F169^, and SSP2^S169^ DAP-seq peaks relative to the transcriptional start (TSS) and end (TES) site of nearby genes (n=8’395). **g-h**, Browser view of SSP, SSP2^F169^, and SSP2^S169^ DAP-seq peaks at *SlGIGANTEA-LIKE1* (g) and *SlGIGANTEA-LIKE2* (h). Normalized coverage (CPM) is shown in yellow, green and blue. Significant peak regions are indicated by red boxes.

### *SSP2* acts redundant with *SSP* to regulate plant architecture

To genetically explore the function of *SSP2*, we used CRISPR-Cas9 genome editing and generated *ssp2^CR^* and *ssp^CR^* null mutants in two determinate cultivars (**Supplementary Fig. 6a-b**). The *ssp^CR^* mutants flowered later than the WT and developed indeterminate shoots, which confirmed previous findings that *SSP* promotes the floral transition (**Fig. 4a-c** and **Supplementary 6c-d**)^15^. We did not observe obvious differences in flowering time for *ssp2^CR^* single mutants, which supports a diminished activity of *SSP2^F169^* in domesticated tomato (**Fig. 4a, c** and **Supplementary 6c-d**). However, *ssp^CR^ssp2^CR^* double mutants tended to flower later than the *ssp^CR^* single mutant, although at high variability (**Fig. 4c and Supplementary 6c, d**). This phenotypic enhancement became more pronounced on sympodial shoots. Double *ssp^CR^ssp2^CR^* mutants produced more leaves on sympodial shoots and more flowers on flowering shoots (inflorescences) (**Fig. 4d, e)**. We concluded that domesticated *SSP2^F169^* is a partial loss-of-function allele and that *SSP* and *SSP2* act partially redundant to promote the transition of meristems to reproductive growth.

**Figure 4:**
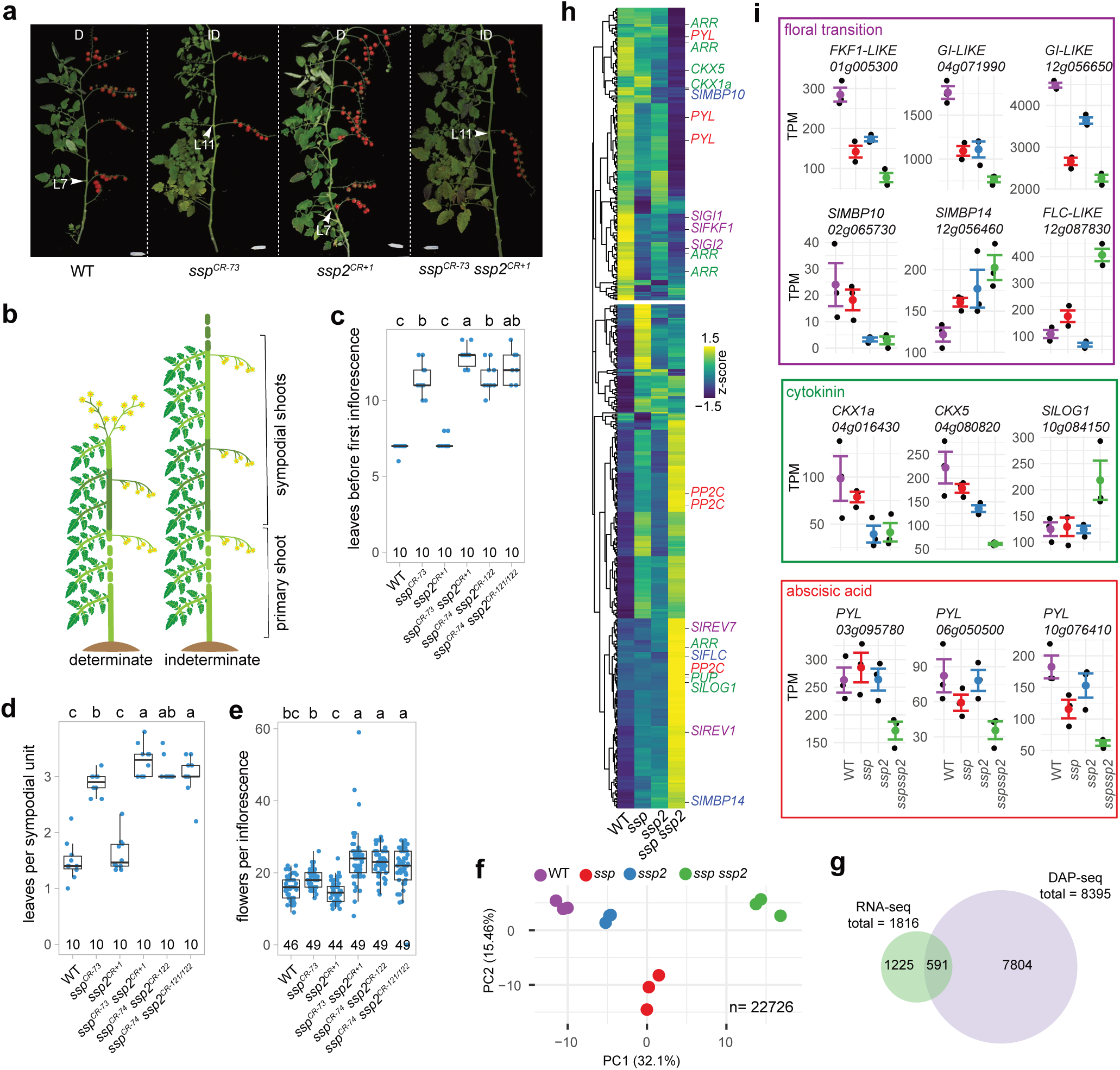
*SSP* and *SSP2* act partially redundant to regulate the transition to flowering. **a**, Representative images of wild-type S100, *ssp^CR^* and *ssp2^CR^* single mutants, and *ssp ssp2^CR^* double mutants. Digits in superscript indicate the number of deleted (CR-) or inserted (CR+) nucleotides. All allele sequences are found in **Supplementary Figure 6**. L= leaf number, arrowheads mark the last leaf before flowering. Determinate (D) and indeterminate (ID) shoots are indicated. Scale bars represent 7.5 cm. **b**, Schematic depiction of tomato shoot architecture. Different shades of green delimit primary and sympodial shoots. **c-e**, Quantification of the floral transition (number of leaves before flowering) on the primary (c) and secondary (d) shoots, and the number of flowers per inflorescence (e) for genotypes shown in (a). The number of plants (c,d) and inflorescences (e) are indicated. Letters represent the results from pairwise comparisons of means using one-way ANOVA and post-hoc Tukey’s HSD test with 95% confidence level. **f**, Principal component analysis of 22’726 expressed genes in transition meristems of the WT, *ssp, ssp2*, and *ssp ssp2*, determined by RNA-seq. **g**, Overlap of genes differentially expressed (log_2_FC ≥ 0.58, FDR ≤ 0.05) in *ssp*, *ssp*2, and/or *ssp ssp2* with genes at SSP, SSP2^F169^, and SSP2^S169^ DAP-seq peaks. **h**, Heatmap depicting expression of 591 putative SSP/SSP2 target genes. **i**, Normalized expression (TPM, transcript per million) for selected putative direct targets in 3 independent biological replicates (black dots) and mean values (colored dots). Error bars indicate standard error. Genes are color coded based on the biological pathway. For box plots in (c-e), the bottom and top of the boxes represent the first and third quartile, respectively, the middle line the median and the whiskers the maximum and minimum values.

To obtain molecular insights into how *SSP* and *SSP2* promote meristem transitions, we sequenced mRNA from micro-dissected meristems at the transition (TM) stage of meristem maturation of the *ssp^CR^* and *ssp2^CR^* single and double mutants, and the WT (in cv. M82)^15^. Clustering of samples in a principal component analysis (PCA) was consistent with the mutant phenotypes that indicated a delayed transition of *ssp^CR^ssp2^CR^*double mutants compared to the *ssp^CR^* single mutant (**Fig. 4f**). We identified 1,816 differentially expressed genes (DEGs) that changed in expression by more than 1.5-fold in at least one of the mutants compared to the WT (FDR ≤ 0.05) (**Supplementary Fig. 6e-f**). Of those, 591 (32.5%) were nearby DAP-seq peaks, indicating that they are direct targets of SSP and/or SSP2 **(Fig. 4g)**. Clustering of the 591 putative direct targets revealed two main patterns of gene expression that contained genes either down-or upregulated (de-repressed) in the *ssp^CR^ssp2^CR^* double mutant (**Fig. 4h,i and Supplementary Table 6**). Among the downregulated genes, we found both tomato homologs of the Arabidopsis floral promoter *GI*, and a homolog of its interactor *FLAVIN-BINDING, KELCH REPEAT, F-BOX 1* (*FKF1*)^29^. In addition, the MADS-box gene *SlMBP10*, a homolog of the Arabidopsis floral promoter *FUL*, was downregulated in *ssp^CR^ssp2^CR^*double mutants, while *SlMBP14* and a *FLOWERING LOCUS C* (*FLC*)-like gene were de-repressed in *ssp^CR^ssp2^CR^*. The MADS-box genes *SlFUL1* and *SlFUL2*, which we used for the initial reporter assays (**Fig. 2f**), were not among the direct targets although both genes were significantly downregulated in the *ssp^CR^ssp2^CR^* double mutant (**Supplementary Fig. 6g**), suggesting that transactivation in the heterologous tobacco system was indirect. We also identified several putative direct targets involved in phytohormone signaling. Two cytokinin dehydrogenase/oxidase genes (*CKX1a*, *CKX5*) and putative negative regulators of cytokinin levels were downregulated while a cytokinin activating enzyme encoding *SlLONELY GUY1* (*SlLOG1*) gene was de-repressed in *ssp^CR^ssp2^CR^*. Furthermore, three abscisic acid receptor genes (*PYLs*) were downregulated in the *ssp^CR^ssp2^CR^* double mutant. These data indicate that *SSP* and *SSP2* redundantly regulate the expression of central regulators of the floral transition and phytohormone responses, and guide meristem transitions towards floral fate.

We revisited the hypothesis that SSP2^F169^ neofunctionalized and compared the transcriptomes of the WT (*SSP2^F169^*) to *ssp2^CR^*single mutants (*SSP2^F169^* loss-of-function) but found only one (Solyc11g071290, alcohol dehydrogenase) of the 230 private SSP2^F169^ DAP-seq targets **(Supplementary Table 7**) to be differentially expressed (**Supplementary Fig. 5h**). Furthermore, Solyc11g071290 was also differentially expressed in the *ssp^CR^* mutant, indicating that misexpression was not specific to loss of *SSP2^F169^* activity (**Supplementary Fig. 5i**). The finding that loss of *SSP2^F169^* activity does not lead to specific misexpression of private SSP2^F169^ targets in the *ssp2^CR^*mutant suggests that the SSP2^F169^ variant did not neofunctionalize but rather compromised its function at targets that are redundant with SSP.

### *SSP2* was lost during the evolution of *Physalis grisea*

To determine whether genetic redundancy between *SSP* and *SSP2* is evolutionary conserved, we inspected orthologs across eudicots (**Supplementary Fig. 7**). Surprisingly, our phylogenetic analyses indicated that tomato *SSP*/*SSP2* and Arabidopsis *FD*/*FDP* resulted from independent duplication events that occurred after the *Solanacaeae* and *Brassicaceae* lineages diverged (**Supplementary Fig. 7**). When we inspected protein sequences of SSP-like transcription factors in the *Solanaceae*, we identified an additional missense mutation (alanine to serine) in a conserved residue in the DNA-binding domain of the potato (*S. tuberosum*) StSSP ortholog, which occurred after the divergence of tomato and potato (**Fig. 2c)**. Furthermore, we found only one SSP-like ortholog in *Physalis grisea* (*PgSSP;* Phygri02g013770), a relative of tomato in the *Solanoideae* subfamily^30^. Phylogenetic and synteny analyses supported an evolutionary scenario in which the ortholog of *SSP2* was lost in *P. grisea* (**Fig. 5a and Supplementary Fig. 8a-c**). To obtain experimental evidence for the loss of redundancy in *P. grisea*, we mutated *PgSSP* by CRISPR-Cas and quantified effects on shoot architecture (**Fig. 5b**). Wild-type *P. grisea* plants produce seven leaves on the primary shoot before terminating in a single-flowered inflorescence (**Fig. 5c**). As in pepper (*Capsicum spp*)^31^, growth continues from two sympodial meristems that are released in the axil of the last two leaves preceding the flower on the primary shoot. Each sympodial meristem gives rise to one sympodial shoot unit, which results in a bifurcation of the shoot. The leaf subtending the sympodial meristem is carried upwards due to strong internode elongation below the sympodial unit and displaced above the preceding flower^31^. Each sympodial meristem produces two leaves and one flower, and in turn releases two additional sympodial shoots. We observed alterations to this pattern in two independent *Pgssp^CR^* mutant lines, which produced an additional sympodial shoot at the first bifurcation and grew less compact than the WT (**Fig. 5c, d-g**). The additional sympodial shoot on *Pgssp^CR^* mutants resulted from a sympodial meristem in the axil of an extra leaf that was produced before flowering, which indicated that loss of *PgSSP* leads to a mild but highly significant delay in flowering time (**Fig. 5e, h**). Together, these results suggest that *PgSSP* regulates the transition of primary and sympodial meristems in the absence of a direct paralog. However, *PgSSP* may act redundantly with other bZIP genes to regulate the floral transition as recently reported in Arabidopsis^32^.

**Figure 5:**
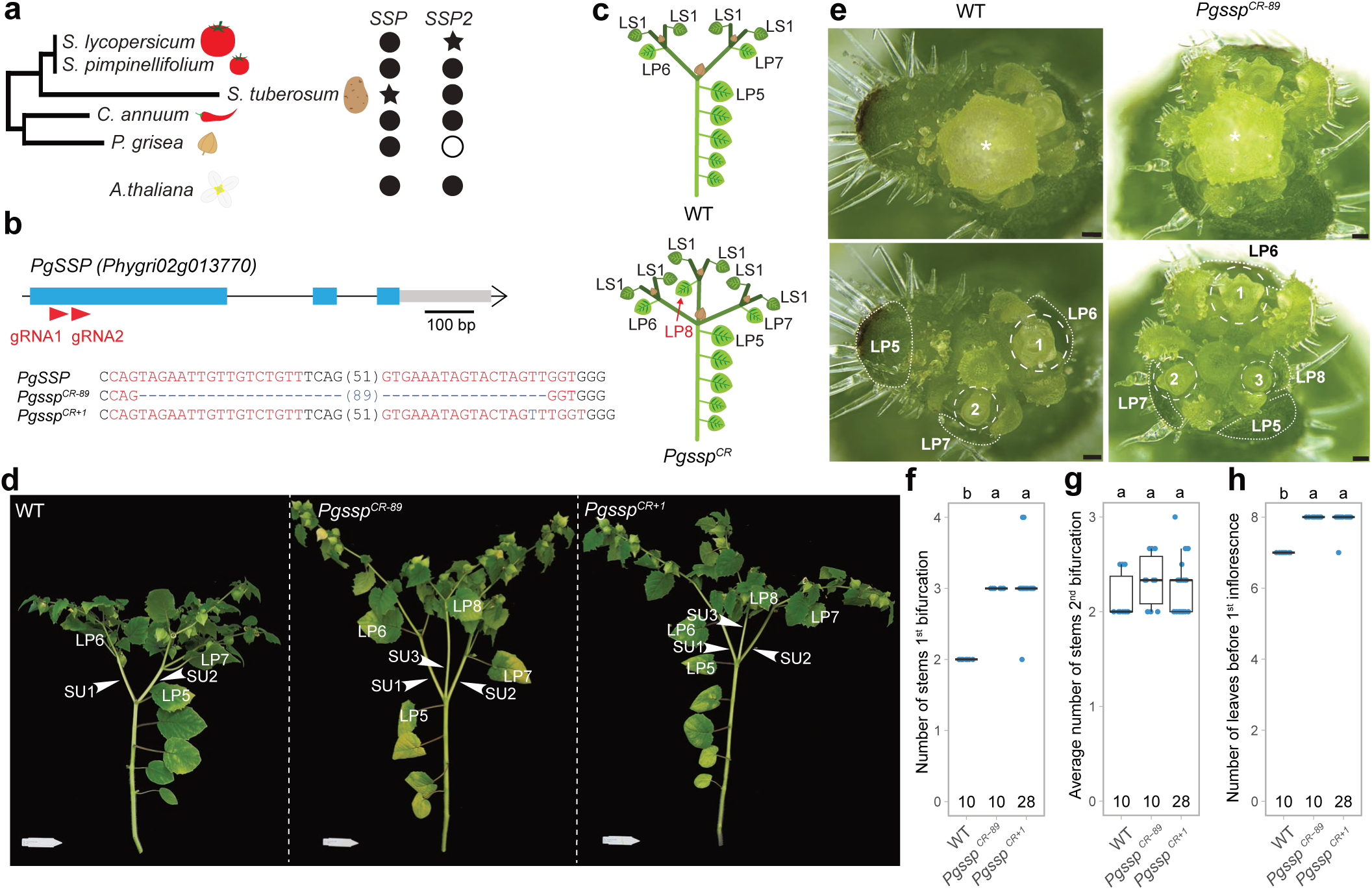
The genome of *Physalis grisea* encodes a single direct *SSP* ortholog that regulates meristem transitions. **a**, Schematic phylogenetic tree of tomato and closely related *Solanaceae* species. Filled circles, empty circles or stars show presence, absence, or missense mutation, respectively, of *SSP*/*SSP2* or *FD/ FDP* in these species. Full tree is displayed in **Supplementary Fig. 7**. **b**, CRISPR-Cas9 targeting of *PgSSP* in *P. grisea*. Blue boxes, black lines, and grey boxes represent exonic, intronic, and untranslated regions, respectively. Single guide RNAs (sgRNAs) are indicated with red arrowheads. PAM and sgRNA sequences are indicated in black and red bold letters, respectively; deletions are indicated with blue dashes; sequence gap length is given in parenthesis. Insertions are indicated by blue letters. **c**, Model of the sympodial growth habit of *P. grisea* WT and *Pgssp^CR^* plants. Different shades of green delimit primary, first sympodial, and second sympodial shoots. The color of leaves on the primary (LP) and sympodial (LS) shoots corresponds with the shoot of origin **d**, Representative pictures illustrating differences in number of sympodial shoots in WT and *Pgssp* mutants. Last leaf before shoot bifurcation (LP5) and displaced leaves (LP6-8) are indicated (L5). White arrowheads indicate individual sympodial units (SU). Scale bar represents 7.5 cm. **e**, Representative stereoscope images of the shoot apex of WT and *Pgssp* mutants. Upper images show the apex with a terminal flower (*). Lower images show the same view with the flower removed. The sympodial meristems (SYMs) and position of removed primary shoot leaves (LP) are delimited by dashed and dotted lines, respectively and numbered in developmental order. Scale bar represents 100 µm. **f-h**, Quantification of the number of sympodial shoots at the first (f) and second (g) bifurcation, and flowering time (number of leaves before the first inflorescence) (h). Number of plants is indicated at the bottom of plots. Letters represent results from pairwise comparisons of means using one-way ANOVA and post-hoc Tukey’s HSD test with 95% confidence level. For box plots in (f-h), the bottom and top of boxes represent the first and third quartile, respectively, the middle line the median and the whiskers the maximum and minimum values.

### Repairing *SSP2* by base-editing leads to earlier fruit yield

Our findings in tomato show that *SSP2* acts partially redundant with *SSP* to promote the transition to flowering on sympodial shoots (**Fig. 4a, d**). We asked whether restoring the activity of *SSP2* in domesticated tomato by correcting the deleterious variant would accelerate the floral transition. We tested this hypothesis by repairing the deleterious variant in domesticated tomato by CRISPR-Cas base editing. The critical non-synonymous mutation results from a TCC (Ser) to TTC (Phe) codon exchange (**Fig. 6a**). The correction of this mutation requires a A-to-G transition on the reverse strand, which can be induced with an adenine base editor (ABE)^33^. Since none of the nearby canonical PAMs (NGG) allowed us to position the target nucleotide into the high-activity editing window (A4-A8) of the protospacer, we used the PAM-less Cas9 variant SpRY fused to ABE8e (**Fig. 6a**)^34^. We edited *SSP2* in the domesticated and double-determinate S100 background^35^ and observed high editing efficiency with edits at the target adenine in 37.5 % (3 of 8) second-generation (T1) transgenic families (**Fig. 6b** and **Supplementary Fig. 9a**). In one T1 family we also detected editing at the bystander T position (**Supplementary Fig. 9a**). To determine whether the base-edited (be) *ssp2^S169be^* allele affected flowering time and shoot architecture, we generated a segregating (F4) population descending from single, heterozygous F3 individual, and selected homozygous (*ssp2^S169be^/ssp2^S169be^*) and heterozygous (*ssp2^S169be^*/*SSP^F169^*) individuals for the repaired allele, and WT siblings (*SSP^F169^*/*SSP^F169^*) as controls by genotyping (**Fig. 6c**). We found that plants homozygous or heterozygous for the repaired *ssp2^S169be^* allele did not flower earlier than their WT siblings (**Fig. 6d-e**). However, they developed less sympodial shoot units and less flowers per inflorescence compared to WT siblings homozygous for the domesticated (*SSP2^F169^*) allele, which resulted in an overall more compact architecture (**Fig. 6d, f-g**). To assess if repair of *SSP2* could compensate for the loss of *SSP*, we introduced the repaired *ssp2^S169be^* allele into the *ssp^CR^*null mutant (in cv. S100). We found that *ssp2^S169be^* did not suppress late flowering and indeterminate growth of *ssp^CR^* (**Fig. 6h-j**). However, we observed a partial and significant suppression of late flowering on sympodial shoots (**Fig. 6h, k**). Moreover, *ssp^CR^ ssp2^S169be^* plants developed shorter inflorescences compared to *ssp^CR^* mutants and WT (*SSP2^F169^*) plants (**Fig. 6l**). Together, these results demonstrate that functional *SSP2* accelerates the reproductive transition of meristems on sympodial shoots in partial redundancy with *SSP*.

**Figure 6:**
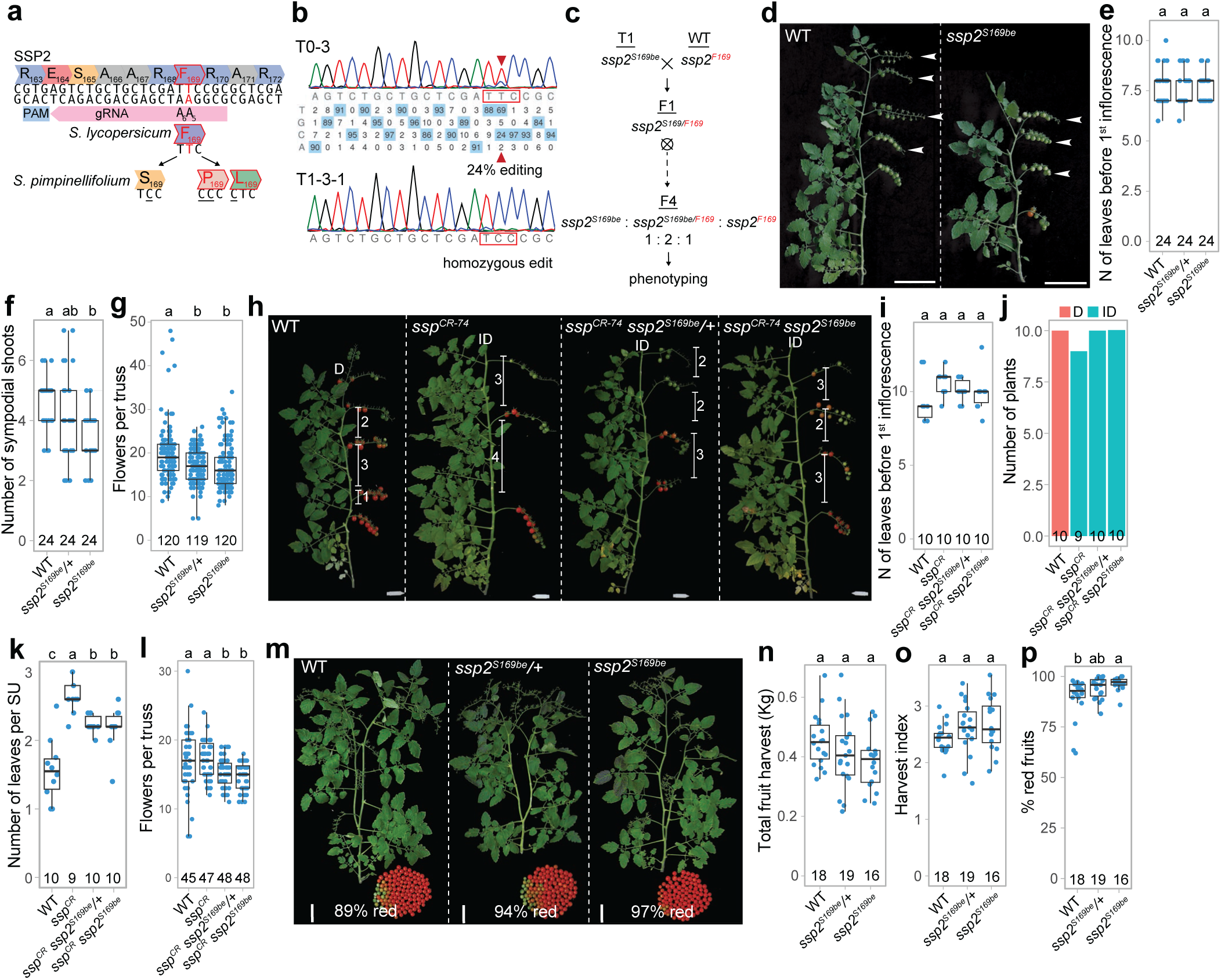
Repairing the deleterious *SSP2* mutation in domesticated tomato by base-editing leads to compact growth and earliness for yield. **a**, Base-editing strategy to correct the *SSP2* mutation using an adenosine base editor (ABE) and a PAM-less Cas9 variant. The target adenine in *SSP2* (A5) is at position 5 of the protospacer with a bystander adenine (A6) at position 6. Editing of the target codon (TTC) can lead to three different outcomes depending on the deaminated adenine. **b**, Validation of editing in a chimeric first-generation (T0) transgenic and corresponding T1 progeny by Sanger sequencing. The target nucleotide is marked by a red arrowhead. **c**, Crossing scheme to generate the segregating *ssp2^S169be^* F4 population. **d**, Representative pictures showing the number of sympodial units on WT and *ssp2^S169be^* plants. Terminal inflorescences of sympodial units are indicated by white arrowheads. **e-g**, Quantification of the number of leaves before the first inflorescence (flowering time) (e), sympodial shoots (f), and flowers per truss (g) of WT, *ssp2^S169be^/+* and *ssp2^S169be^* plants. **h**, Representative pictures showing the number of leaves per sympodial unit and determinacy of WT, *ssp^CR^*, *ssp^CR^ ssp2^S169be^/+* and *ssp^CR^ ssp2^S169be^* plants. **i-l**, Quantification of flowering time (i) (as in (e)), number of determinate plants (j), leaves per sympodial unit (SU) (k), and flowers per truss (l) of WT, *ssp^CR^*, *ssp^CR^ ssp2^S169be^/+* and *ssp^CR^ ssp2^S169be^* plants. Determinate (D) and indeterminate (ID) shoots are indicated. **m**, Representative images showing the complete harvest of individual WT, *ssp2^S169be^/+* and *ssp2^S169be^* plants. Percentage of red fruits is indicated. **n-p**, Quantification of total fruit yield (n), harvest index (total fruit yield / plant weight) (o), and percentage of red fruits. Number of plants are indicated in the plots for (e-g), (i-k) and (l-o). Letters on top of the plots represent pairwise comparisons of means using one-way ANOVA and post-hoc Tukey’s HSD test with 95% confidence level. Scale bars represent 10 cm (d) and 7.5 cm in (h,m). For box plots in (e-g, i, k, l, n-p), the bottom and top of boxes represent the first and third quartile, respectively, the middle line the median and the whiskers the maximum and minimum values.

Tomato production was revolutionized during the 20^th^ century by the *self-pruning* mutation, which confers determinate growth and facilitates mechanical harvesting. Our findings showed that a functional *SSP2* allele accelerates sympodial shoot flowering and thus suggested an agronomic value for this allele regarding earliness for yield. To test whether accelerated flowering from the repaired *ssp2^S169be^* allele leads to earlier yield, we quantified fruit production in segregating (F4) populations under experimental greenhouse conditions (see **Online Methods**). We found that total fruit yields, harvest index, and fruit size for *ssp2^S169be^* plants were comparable to the WT sibling controls (**Fig. 6n-o** and **Supplementary Fig. 9b-e**). Notably, *ssp2^S169be^*homozygotes displayed an 8% increase in the proportion of ripe fruits compared to WT siblings, which was likely due to precocious flowering and termination of sympodial shoots (**Fig. 6m, p**). Thus, compact growth from repairing the deleterious *SSP2* mutation by base editing can confer earliness for fruit yields and represents a new target for customizing tomato shoot architecture. However, *ssp2^S169be^* fruits had a reduced sugar content (brix) (by 11%) (**Supplementary Fig. 9f**), suggesting trade-offs between compact growth with soluble sugar content of fruits.

## DISCUSSION

Here, we investigated the load of deleterious mutations that accumulated during domestication and improvement of tomato. Within genes central to flowering time control, we discovered a deleterious variant in the previously uncharacterized bZIP transcription factor gene *SSP2*. The deleterious variant results in the exchange of a conserved serine to a phenylalanine in the DNA-binding domain of the transcription factor. Our results from structural modelling, genome-wide DNA binding assays, and genetic analyses indicate that the domesticated SSP2^F169^ variant partially lost its ability to bind and regulate target genes that are largely shared between the ancestral SSP2^S169^ variant and its paralog SSP. However, we cannot fully rule out that domesticated SSP2^F169^ neonfunctionalized given its 230 private targets genes and ability to bind both a G-box and an alternative TA-rich target motif, although at low affinity. Interestingly, in the yeast bZIP factor Pap1, the equivalent serine-to-phenylalanine exchange contributes to a similar change in binding specificity for <u>TTA</u>CG<u>TAA</u>/<u>TTA</u>G<u>TAA</u> sequences^36^. However, we found only very few target genes with this alternative motif bound by SSP2^F169^ and none of them was specifically differentially expressed in the *ssp2^CR^* mutant, in which SSP2^F169^ activity is lost. We conclude from this data that the deleterious variant in *SSP2* led to loss of genetic redundancy between *SSP* and *SSP2*, a pair of paralogs that is widely conserved in flowering plants. In Arabidopsis, it was shown that *FD* and *FDP* act redundantly during phytohormone responses while only *FD* affects the floral transition, suggesting functional divergence of *FD*^21^. In contrast, our findings in tomato indicate that *SSP* and *SSP2* act also partially redundant during the floral transition. Notably, our phylogenetic analyses suggest that paralogs of *SSP* and *FD* arose independently in *Solanaceae* and *Brassicaceae*, which could explain species-specific divergence of this paralogous pair. The complete loss of a *PgSSP* paralog in *Physalis grisea* further supports dynamic evolution of the paralog pair. Deleterious mutations and gene loss have been proposed as an important mechanism of adaptation^6,7^. However, the benefit of the deleterious *SSP2^F169^* variant during domestication remains speculative. Our genetic data demonstrates that domesticated *SSP2^F169^* delays meristem transitions on shoots and inflorescences. Notably, the domesticated *SSP2^F169^* genotype develops more flowers per inflorescence than the ancestral *SSP2^S169^* genotype. Although flower number correlates with fruit yield, the number of flowers per inflorescence in general decreased during tomato domestication, likely due to source-sink imbalances driven by dramatic increases in fruit size^37^. This overall decrease in flower number during tomato domestication suggests that effects from *SSP2^F169^* on flower number were rather minor and difficult to select. Furthermore, the deleterious *SSP2^F169^* variant could have hitchhiked near QTLs that were selected during domestication and improvement, which is a common scenario in crops with a narrow genetic base such as tomato^38^. However, the closest known improvement sweep on chromosome 2 with five fruit-weight QTLs is more than 5 Mbp away from *SSP2*, rendering linkage unlikely^38^. Finally, we cannot exclude that *SSP2^F169^* is adaptive under specific conditions or developmental processes that were absent from our experiments given that *CETS* genes play roles beyond flowering time control^39^.

Whether *SSP2^F169^* was nearly fixed in cultivated tomato due to selection or drift remains therefore an open question. Yet, the loss of genetic redundancy caused by a deleterious mutation may reflect a common feature during the selection of crops in human-made environments. The less-is-more idea proposes the accumulation of loss-of-function mutations as a driver of rapid evolutionary change^7^, and gene loss may be even more frequent during the intense artificial selection in domesticated environments. A reduced genetic repertoire in domesticated genomes could result in lower genetic redundancy compared to their ancestral states and, as a consequence, facilitate the exposure and selection of novel mutations, which are otherwise masked by redundant paralogs. Our data shows that the ancestral *SSP2^S169^* allele can suppress effects of *ssp* mutations, which allow tuning of shoot architecture and optimization of tomato yields^15^. Intriguingly, the deleterious *SSP2^F169^* mutation, which broke redundancy with the paralog *SSP*, may have been a prerequisite for the identification of the *ssp^2129^* breeding mutation. This illustrates how standing variants can become adaptive due to genetic interactions with mutations that are introduced or arose during breeding.

Correcting deleterious variants with genome editing in crops has been proposed as major strategy for future crop breeding^4^. To our knowledge, we present here the first example of a direct repair of a deleterious mutation in a crop using base editing. We show that repairing the deleterious *SSP2* variant in tomato leads to precocious flowering on sympodial shoots and an overall more compact plant architecture, although with a negative consequence on fruit sugar content. Notably, precocious flowering and compact growth of base-edited plants were associated with earliness for yield, with repaired plants displaying an 8% increase in ripe fruits at harvest. Such earliness for fruit yield is a highly desirable trait for customizing shoot architecture for specific environments. Our work shows that base editing provides a promising approach for correcting deleterious variants that accumulated during domestication and improvement in crops. However, our study also emphasizes that deleterious mutations are not unfavorable *per se* and may have adaptive roles that are only exposed in specific genetic backgrounds or environmental conditions.

## ACKNOWLEDGEMENTS

We thank all members of the Soyk lab, Y. Eshed, and C. Fankhauser for helpful discussions; J. Marquis and J. Weber for support with sequencing; B. Tissot, L. Nerny, V. Vashanthakumar, Y. Emmenegger, A. Chatillon, L. Keel, and T. Stupp for support with plant care; G. Ghazi Soltani and S. Mainiero for support with experiments; J. M. Franco-Zorrilla for advice with DAP-seq; J. van Eck and K. Swartwood for advice with plant transformation; Z. Lippman, Y. Qi, and T. Jacobs for providing materials. This work was supported by the University of Lausanne, the European Research Council (ERC) under the European Union’s Horizon 2020 research and innovation programme (ERC Starting Grant “EPICROP” Grant No. 802008) to S.So., the Swiss National Science Foundation (SNSF) under an Eccellenza Professorial Fellowship (Grant No. PCEFP3_181238) and Project Grant (Grant No. 310030_212218) to S.So., and an UNIL Interdisciplinary Project Grant to N.Gl. and S.So., and an National Science Foundation Grant (IOS-1546625) to G.B.M and S.St..

## AUTHOR CONTRIBUTIONS

A.N.G., S.St., and S.So. conceived the project and designed and planned experiments

A.N.G., M.B, G.S., L.L., J.I., I.J., J.Z., S.So. performed experiments and collected data

A.N.G., M.B, J.I., G.A., I.J., J.Z., R.R., C.I., N.Gu., J.J.-G., N.Gl., S.St., S.So. analysed data

N.Gl., G.B.M., S.St., S.So. aquired project funding.

A.N.G. and S.So. wrote the first draft of the manuscript

All authors read, edited, and approved the manuscript.

## ONLINE METHODS

### Plant material, growth conditions, and phenotyping

Seeds of *S. lycopersicum* cv. M82 (LA3475), *S. lycopersicum* cv. Sweet-100 (S100) double-determinate^35^, *S. pimpinellifolium* (LA1589), *P. grisea,* and *N. benthamiana* were from our own stocks. Tomato seeds were germinated on soil in 96-cell plastic flats. *P.grisea* seeds were incubated at 50°C for 3 days prior to sowing to increase germination rates. Plants were grown under long-day conditions (16-h light/ 8-h dark) in a greenhouse supplemented with artificial light from high-pressure sodium bulbs (∼250umol m-2s-1). Temperature was 25°C and relative humidity was 50-60%. Tomato plants were grown in 5L pots (2 plants per pot) and physalis in 0.36 l pots (one plant per pot) under drip irrigation and standard fertilizer regimes. Tomato plants were pruned down to the primary and proximal axillary shoots. Phenotypic data was collected from the F3 and T4 generation for *ssp^CR^ ssp2^CR^* plants in the S100 background, the F7 (*ssp^CR^*and *ssp2^CR^*) and F4 (*ssp^CR^ ssp2^CR^*) generation in the M82 background, and the T3 generation for *PgSSP^CR^*in *Physalis* and the F4 generation in *ssp ssp2^S169be^* plants. Data for flowering time, sympodial shoot number, number of leaves per sympodial shoot, and number of flowers per inflorescence were collected from primary and proximal shoots. To assess tomato yield components under experimental greenhouse conditions, mature plants were harvested 79 days after transplanting. For data collection, plants and fruits were manually removed from the soil and the plant, respectively. Total fruit yield was defined as the sum of red and green fruits from each plant. Harvest index was calculated by dividing total fruit yield by plant weight (i.e., vegetative biomass after fruit removal). Ten fruits per plant were randomly selected to measure average fruit weight and total soluble sugar content (brix). Brix was quantified using a digital Brix refractometer (HANNA® instruments, HI96801). Statistical analyses of phenotyping data were conducted in R^40^.

*N. benthamiana* (tobacco) seeds were sown on soil in square pots. Seedlings were grown under long-day conditions (16-h light/ 8-h dark) in a growth room under LED light (∼120 umol m-2s-1) and constant temperature (25°C). Approximately one week after germination, tobacco seedlings were singled out into square pots.

### LA1589 de novo genome assembly

Nanopore long read sequences for the *S. pimpinellifolium* accession LA1589 were previously generated^16,41^. Basecalling was performed using Guppy v3.1.5. Illumina sequencing data were previously generated^24^. We assembled the Nanopore and Illumina sequences together with MaSuRCA (v3.4.1)^42^. Resulting contigs were scaffolded against the Heinz 4.0 reference genome using RaGOO (v1.1)^43^. Gaps were closed with LR_Gapcloser (v3)^44^ and the assembly was polished with 3 rounds of Pilon (v1.23)^45^. Assembly statistics are found in **Supplementary Table 8**. We used liftoff^46^ to annotate the LA1589 assembly with ITAG4.0 gene models and tomato pan-genome genes as previously described^35^.

### Genome-wide prediction of deleterious variants

Illumina raw reads from 27 *S. pimpinellifolium*, 23 *S. lycopersicum* var. *cerasiforme*, and 32 *S. lycopersicum* accessions (**Supplementary Table 1**) were retrieved from public repositories as described before^47^. Reads were aligned to the *S. pimpinellifolium* reference genome (LA1589v0.1) using BWA-MEM (v0.7.17) using default parameters. Alignments were sorted and duplicates marked with PicardTools (v2.26.2) and indexed using samtools (v1.15.1)^48^. Variants were called with bcftools (v.1.15.1, parameters mpileup --no-BAQ --ignore-RG -d 1000000 -Q0 --annotate FORMAT/AD,FORMAT/DP). Variants were filtered with vcftools (v0.1.14, parameters --min-alleles 2 --max-alleles 2 --minQ 30 --minDP 5 --maxDP 50 --mac 2 --recode --recode-INFO-all). Filtered variant call format (vcf) files were used to predict deleterious mutations using SIFT-4G^17^. A SIFT database was built from the *S. pimpinellifolium* genome sequence (SpimLA1589_v0.1) and annotation (SolpimLA1589_v0.2) using the SIFT instructions and default parameters. The LA1589 SIFT database contained SIFT scores for 70% of genes (21578 of 30808), SIFT scores for 83% of positions (56424493/67919880), and confident scores for 73% of positions (41083097/56424493). SIFT was used to determine the effect of coding sequence variants on protein sequence, and to predict deleterious missense variants. Variant types and SIFT scores were plotted in R using the ggplot2 package.

### Phylogenetic analyses and sequence alignments

Protein sequences of tomato and Arabidopsis bZIP family members were obtained from the Plant Transcription Factor Database (PlantTFDB, v5.0)^49^. Physalis bZIP protein sequences were identified in a BLAST search on the Phygri1.3.1 protein annotation^30^ using SSP protein as query. Full-length sequences of 70 tomato, 74 Arabidopsis, 58 Physalis, and yeast Pap1 (SPAC1783.07c.1) bZIP proteins were aligned with MAFFT (v7.481) using default parameters^50^. Maximum likelihood phylogenetic trees were constructed in IQ-Tree (v2.2.0.5; parameters -m MFP -bb 1000 -bnni -redo)^51^ and visualized in FigTree (v1.4.4; http://tree.bio.ed.ac.uk/software/figtree/). Average number of substitutions per site is indicated by scale bars. Specific bZIP groups were assigned according to their Arabidopsis homologs^52^.

To reconstruct the phylogenetic tree of the bZIP family in eudicots we used the OMA browser’s^53^ 07/2023 release to collect homologs for tree building. Hierarchical Orthologous Groups (HOGs) were identified by searching for the SSP gene identifier (Solyc02g083520) for the initial HOG and adding additional closely related HOGs, inferred to be closely related as they share many predicted orthologs. The following HOGs were downloaded: D0228852, D0178917, D0181214, D0210160, D0214417, D0216285, D0223413 (accessed 23 Jan 2024). Additionally, through BLAST searches, we incorporated the bZIP gene of *Amborella trichopoda* and closely related bZIP genes from eight *Solanaceae* species: *Nicotiana benthamiana*, *Nicotiana tabacum*, *Physalis grisea*, *Petunia axillaris*, *Petunia inflata*, *Solanum tuberosum*, *Capsicum annuum*, and *Capsicum chinense*. The final dataset comprised 128 genes from 51 plant species. These protein sequences were aligned using the approach described in the PhylomeDB pipeline^54^. Briefly, we obtained alignments in forward and reverse directions using three programs (MUSCLE v3.8.1551^55^, MAFFT v7.490^50^, and Kalign v3.3.5^56^). Then, the six alignments were combined using M-COFFEE v13.46.0.919e8c6b^57^. The phylogenetic tree was reconstructed using a maximum likelihood approach in IQ-TREE v2.2.2.6^58^, using the best-fit model identified by ModelFinder^59^ (JTT+F+I+R5) and 1000 ultrafast bootstrap replicates. The tree was manually rooted using *Amborella trichopoda* as outgroup. Duplication events were inferred using ETE v4.0^60^ and the species overlap method^61^.

### Homology modelling

The HHpred server was used to find suitable templates for SSP2 protein modeling^62^. The final templates were chosen based on sequence similarity in the area of protein-DNA interaction. The 50 homology models of wild tomato protein SSP2^S169^ dimers were calculated using Modeller 9v18^25^ and CCAAT/enhancer-binding protein beta (C/EBP beta) as a template. The crystal structure of human C/EBP beta in complex with DNA is stored under 1HJB code in the Protein Data Bank^26^. The target and template sequence shared 26% of sequence identity. The best model in terms of its DOPE score^63^ was chosen. Analogically, the 50 homology models of domestic tomato SSP2^F169^ protein dimers were calculated based on the structure of Pap1 as a template and the best model, according to DOPE score, was chosen. The crystal structure of Pap1 is stored in the PDB under 1GD2 code and shares 24% of sequence identity with the SSP2^F169^ protein^36^. For both SSP2 proteins the DNA molecule from the template structure was included in the models. The DNA sequence was changed to the SSP2 recognition motif with UCSF Chimera tool that was also used for visualization of the models^64^.

### Molecular cloning

Binary vectors for CRISPR-Cas9 mutagenesis in tomato were assembled using the Golden Gate cloning system as previously described^35,65^. For CRISPR-Cas9 mutagenesis, a new Level (L) 1 part pICH47742_SpCas9-P2A-GFP was cloned by amplifying SpCas9 from pICH47742::35S::Cas9 (Addgene no. 49771) using primers P94 and P129. The fragments were cloned into the L0 acceptor pAGM1287 to generate pAGM1287-SpCas9. P2A-GFP was amplified from pGG-D-P2A-GFP-NLS-E^66^ using primer P96 and P97 and cloned into the L0 acceptor pAGM1301 to generate pAGM1301_P2A-GFP. The pAGM1287_SpCas9 and pAGM1301_P2A-GFP parts were combined with pICH51288 (2Xp35S) and pICH41421 (nosT) in pICH47742 (L1 acceptor) to generate pICH47742_SpCas9-P2A-GFP. For CRISPR-Cas base editing, the PAM-less adenosine base editor ABE8e-SpRY^34^ was domesticated by amplifying four fragments using the primer pairs P576/ P577, P578/ P579, P580/ P581, P582/P583 on the template pYPQ262B^34^. Fragments were cloned into the L-1 acceptor pAGM1311 and combined in the L0 acceptor pAGM1287 to generate pAGM1287_ABE8e-SpRY. pAGM1287_ABE8e-SpRY was combined with pAGM1301_P2A-GFP, pICH51288 (2Xp35S), and pICH41421 (nosT) in the L1 acceptor pICH47742 to generate pICH47742_SpRY-ABE8e-P2A-GFP. Constructs for transactivation assays were cloned using the MoClo kit^65^. The p19 construct for silencing suppression was assembled with the L1 acceptor pICH47742 and L0 parts pICH85281 (pMas), pICH44022 (p19), and pICH77901 (tMas). The YFP construct was assembled with the L1 acceptor pICH47742 and L0 parts pICH51266 (p35S), pICSL80014 (YFP), and pICH41414 (t35S). To clone the SFT co-effector and the SlycSSP2 effector constructs, the coding sequences of SFT and SlycSSP2 were amplified from *S. lycopersicum* (cv. M82) transition meristem cDNA with gene specific primer pairs (SFT: SFT_F/SFT_R, SlycSSP2: SSP2_F/SSP2_R). To clone the SpimSSP2 effector construct, the coding sequence of SpimSSP2 was amplified from *S. pimpinellifolium* (LA1589) transition meristem cDNA with the primer pair SSP2_F/SSP2_R. The amplicons were cloned into the L0 acceptor pICH41308. Individual L0 effector parts (SlycSSP2, and SpimSSP2) were combined with pICSL13001 (p35S), pICSL30009 (Myc-tag), and pICH41414 (t35S) in the L1 acceptor pICH47772. The L0 co-effector part (SFT) was combined with pICSL13001 (p35S), pICSL30008 (HA-tag) and pICH41414 (t35S) in the L1 acceptor pICH47761. To clone the SSP effector construct and the 35S::YFP no-effector construct the coding sequence of SSP was amplified from *S. lycopersicum* (cv. M82) transition meristem cDNA with the primer pair SSP_F/SSP_R and the coding sequence of 35S::YFP was amplified from pENTR_L1-Turbo-YFP-NLS-STOP-L2 (addgene no. 127361) with the primer pair YFP_F/YFP_R. Both amplified coding sequences were cloned into the L1 acceptor pICH47772 that was amplified with the primer pair TEA_F/TEA_R with the NEBuilder HiFi DNA Assembly Cloning Kit (NEB #E5520). To clone the luciferase reporter constructs, upstream regions of pMC, pFUL, and pFUL2 were amplified from *S. lycopersicum* (cv. M82) gDNA in multiple fragments gene-specific primer pairs (pMC: pMC_F1/pMC_R1 and pMC_F2/pMC_R2; pFUL1: pFUL1_F3/pFUL1_R3 pFUL1_F2p/FUL1_R2p, and FUL1_F1/pFUL1_R1; pFUL2: pFUL2_F1/pFUL2_R1 and pFUL2_F2/pFUL2_R2) and cloned into the L-1 acceptor pAGM1311. The pMC construct contained 2170 bp genomic sequence including upstream region, the 5’UTR, and the first exon. The pFUL1 and pFUL2 constructs contained 2640 bp and 2040 bp genomic sequence, respectively, including upstream regions and the 5’UTR. The L-1 parts were cloned into the L0 acceptor pICH41295. Individual L0 effector parts (pMC, pFUL1, and pFUL2) were combined with pICSL80001 (fLuc) and pICH41432 (tOCS) in the L1 acceptor pICH47751. All primers and gRNA sequences are listed in **Supplementary Table 9 and 10.**

### CRISPR/Cas9 genome editing, plant transformation and identification of mutant alleles

CRISPR-Cas9 mutagenesis in tomato and physalis was performed as described previously^35,67,68^. Briefly, guide RNAs (gRNAs) were designed using the CRISPOR tool and the M82v1.0, Sweet-100v2.0 or Phygriv1.0 genome assemblies. Final vectors were transformed into the tomato cultivar M82, LA1589 or double-determinate Sweet-100, or into *P. grisea* by A*grobacterium tumefaciens*-mediated transformation. CRISPR-Cas9 editing in tomato and physalis was verified by genotyping or amplicon sequencing as described^35^. Base editing was quantified in first-generation (T0) transgenics using EditR v1.0.10^69^ and in the T1 generation with a CAPS marker (**Supplementary Table 9)**.

### Generation of near-isogenic lines (NILs)

Near-isogenic *SSP2^S169^* lines in the M82 background were generated by crossing the *S. pimpinellifolium* accession LA1589 with *S. lycopersicum* cv. M82, and backcrossing F2 individuals homozygous for *SSP2^S169^* to the recurrent parent (*S. lyc.* cv. M82) over 4 (BC4) to 5 (BC5) generations. Presence *SSP2^S169^* allele was confirmed by CAPS marker (**Supplementary Table 9**).

### Transactivation assays

Transient transactivation assays with luciferase reporter constructs were conducted in *N. benthamiana* leaves as previously described^70^. In brief, leaves of 34 and 36 days old plants were infiltrated with mixtures of *A. tumefaciens* (strain GV3101) cultures containing effector, co-effector, luciferase reporter, transfection control, and silencing inhibitor vectors. Effector constructs contained the coding sequence (CDS) of *SSP*, *SSP2^F169^* or *SSP2^S169^* with an N-terminal Myc tag and driven by the CaMV 35S promoter. The co-effector construct contained the CDS of *SFT* with an N-terminal HA tag and driven by CMV 35S promoter. The luciferase reporter constructs contained the CDS of fLUC driven by the upstream regions of *MC*, *SlFUL*, or *SlFUL2*. The transfection control was pGREENII-0800-LUC, which contains the CDS of rLUC driven by the CMV 35S promoter. A 35S::YFP construct was used as no effector control. A p19 construct was used to suppress silencing. Liquid cultures were grown in 10 ml YEB in 50 mL Falcon tubes for 20 hrs at 28°C, shaking. Agrobacteria were harvested by centrifugation at 1932 x g and resuspended in infiltration buffer (50 mM MES pH 5.7 and 10 mM MgCl_2,_ 200 µM Acetosyringone) to an OD_600_ = 0.5. Before leaf infiltration, individual cultures were incubated 3 hrs at RT and combined to obtain mixtures with effectors, reporters (fLUC), and transfection control (pGREEN 35S:rLUC), and silencing inhibitor (p19) plasmids at final OD_600_ of 0.1, 0.1, 0.1, and 0.05. Agrobacteria mixtures were infiltrated into the 4^th^, 5^th^ and 6^th^leaf using a needleless syringe, with three different plants being infiltrated for each combination. Leaf disks of 0.8 cm diameter were harvested 3 days after infiltration and flash-frozen in liquid nitrogen before grinding in a mix mill (twice 20 s^−1^ for 30s). Luciferase assays were performed using the Dual-Luciferase Reporter Assay System (Promega) and a Hidex Sense Microplate Reader. In short, leaf powder was extracted in 300 µl of 1x PLB and vigorously vortexed for 30 s. Volumes of 10 µl protein extracts were mixed with 40 µl luciferase reagent in 96-well microplates and incubated for 10 min at RT. Firefly luciferase (fLUC) activity was quantified (9 s integration time). Reactions were mixed with 25 µl Stop & Glo. Renilla luciferase (rLUC) activity was measured (9 s integration time). Transactivation by the effectors was determined by calculating the fLUC/rLUC ratios.

### DAP-seq

Myc-tagged coding sequences of *SSP*, *SSP2^F169^* and *SSP2^S169^* were amplified from effector constructs used in the transactivation assay. The pTnT™ vector, and the *SSP2^F169^* and *SSP2^S169^* inserts were digested using XhoI (NEB) and NotI-HF (NEB) and combined using T4 Ligase (NEB). The Myc-tagged coding sequence for *SSP* was cloned into pTnT™ vectors with the NEBuilder HiFi DNA Assembly Cloning Kit (NEB #E5520). Plasmid DNA was isolated from 100 ml bacterial cultures using the PureYield™ Plasmid Midiprep System (Promega, A2492). Two replicates of SSP and SSP2 proteins were expressed *in-vitro* in the TnT® SP6 High-Yield Wheat Germ Protein Expression System (Promega, L3260) from 3.5 μg plasmid DNA per reaction. Protein expression and solubility was verified by western blotting using primary anti-Myc (Merck, clone 4A6, 1:5000) and secondary anti-mouse (Amersham ECL NA931, 1:10,000) antibodies, according to the manufacturer’s instructions. High molecular weight DNA for genomic library construction was isolated from inflorescence meristem tissue of the *anantha* mutant in Sweet-100 using a CTAB protocol as described previously^35^. DAP-seq was performed as previously described with minor modifications^27,71^. The DNA-library was prepared according to Franco-Zorilla & Prat (2021) with minor modifications. The gDNA library was purified using SPRI beads (B23317, Beckman Coulter). Adaptor ligation was verified by qPCR with index-specific primers (**Supplementary Table 9**) and KAPA standards 20, 2 and 0.2 nM (Roche) in 10 μl reactions. DNA affinity-purification steps were performed according to Bartlett et al. (2017) with 75 ng of gDNA input library per replicate. Eluted libraries were single-indexed (**Supplementary Table 9**). Eight uniquely-indexed libraries were produced, two replicate libraries per protein (SSP, SSP2^F169^, SSP2^S169^) and two replicates of the input library as negative control. Indexed libraries were purified individually with the Monarch® PCR&DNA Cleanup Kit (NEB, T1030S). Individual indexed libraries were analyzed on a Fragment Analyzer (Agilent), purified with SPRI beads and pooled at equimolar (10 nM) concentrations. Pooled libraries were sequenced on one Illumina NovaSeq6000 lane at the Genome Technology Facility (GTF) of the University of Lausanne. A total of 753’327’838 PE150 reads (between 64’808’988 and 144’444’123 per sample) were generated. Raw read quality was assessed using FastQC (v0.11.9; http://www.bioinformatics.babraham.ac.uk/projects/fastqc/). Adapter sequences were trimmed with NGmerge (v0.3, parameters -g -d -a)^72^. Reads were aligned to the SollycSweet-100v2.0 reference ^35^ with hisat2 (v2.2.0, default parameters)^73^, and alignments were sorted and indexed using samtools (v1.15.1)^48^.

Peak calling and differential binding (DB) analysis was performed with the Bioconductor csaw package (v1.301)^74^. Peak calling in csaw is based on count-based models, in particular the quasi-likelihood (QL) framework of the edgeR package is used^75^. The counts are modelled using NB distributions that account for overdispersion between biological replicates^76^. Each window can then be tested for significant DB between conditions. We used a window width of 10 bp and an estimated fragment length of 213 bp. Prior to counting, repeats were blacklisted from the genome using the SollycSweet-100v2.0 TE annotation^35^. To filter regions and windows, we used the global enrichment approach of csaw. After variance stabilization, windows with significant differential binding with respect to specific factors of interest in our design matrix were identified using the glmQLFTest function. Bins of 10000 bp were used for global background estimation. The median of the average abundances across all 10000 bp bins was used as the global background coverage estimate. We only retained windows with at least a 4-fold change from the global background coverage. We counted the reads into large bins and normalized with the wrapper function normFactors, which uses trimmed mean of M-values (TMM) method. The false discovery rate (FDR) control was obtained by applying the Benjamini-Hochberg (BH) method. In csaw, the FDR control was applied to all detected DB regions or peaks. Regions were obtained by grouping windows into regions using the csaw mergeWindows function. The minimum distance at which two binding events were treated as separate sites was set to 1000 bp. Significant regions were identified with the csaw makeContrasts function (FDR ≤0.01). Gene-based annotation of differentially-bounds regions was performed using the detailRanges function of csaw (3 Kbp upstream and 2 Kbp downstream of TSS) and annotation file SollycSweet-100_genes_v2.1.1.gff3^35^. BED files with significant regions and BigWig files with normalized read coverage were exported via the *export* function of the rtracklayer package^77^ in R. *De-novo* motif discovery was performed with the 1000 most significant peaks (by FDR) for each sample by analysing genomic sequences from position −100 to +100 relative to the peak center using MEME (v 5.3.3; parameters -dna -mod zoops -nmotifs 3 -minw 6 -maxw 15 -maxsites 1000 -objfun classic -revcomp - markov_order 0)^78^.

Genome-wide distribution of peaks was determined using ChIPSeeker (v1.32.0)^79^ by annotating regions +/− 5 Kbp around the TSS with the function annotatePeak (parameters tssRegion=c(−5000, 5000)). Peak intensity profiles and peak heatmaps were generated using the computeMatrix, plotHeatmap, and plotProfile functions in deepTools^80^. Most-enriched motifs for SSP, SSP2^F169^, and SSP2^S169^ were mapped to the SollycSweet-100v2.0 reference^35^ with FIMO/MEME Suite^78^. Browser shots of peak coverage, peak regions and binding motifs at putative direct targets were generated in jbrowse2^81^.

### Gene Ontology enrichment analyses

Gene Ontology (GO) enrichment analyses for biological processes were performed using a functional annotation database for the ITAG4.0 tomato genome from PLAZA^82^ and the ClusterProfiler package in R^83^ with a pvalueCutoff = 0.05 and pAdjustMethod = BH.

### RNA-seq

Meristem staging, collection, RNA extraction, and library construction for the *ssp^CR-181^* (188 bp deletion allele), *ssp2^CR-122^* (122 bp deletion allele) and *ssp^CR-181^ssp2^CR-122^* mutants, and the WT in the genetic background of cv. M82 was performed as previously described^23^. In brief, seedlings shoot apices were collected at the transition (TM) stage of meristem maturation, and immediately submerged in ice-cold acetone. Shoot apices were manually dissected under a stereoscope and three biological replicates consisting of 14-22 meristems were collected per genotype from individual seedlings. Total RNA was extracted with the Arcturus Pico-Pure RNA Extraction kit (Thermo). We prepared indexed libraries using the TruSeq Stranded mRNA Library Prep kit from Illumina according to the manufacturer’s instructions. Fragment size and concentration were assessed with a Bioanalyzer. Libraries were sequenced on 2 Illumina NovaSeq6000 lanes at the Genome Technology Facility (GTF) of the University of Lausanne. A total of 187’907’134 SE100 reads (between 14’133’226 and 17’789’680 per sample) were generated.

The quality of raw reads was assessed using FastQC (v0.11.9; http://www.bioinformatics.babraham.ac.uk/projects/fastqc/). Raw reads were aligned to the genome reference M82v1.0^35^ using STAR^84^ (v2.7.6a; parameters --runMode alignReads -- outFilterType BySJout --outFilterMultimapNmax 20 --outMultimapperOrder Random -- alignSJoverhangMin 8 --alignSJDBoverhangMin 1 --alignIntronMin 20 –alignIntronMa× 1000000 --alignMatesGapMa× 1000000). Alignments were sorted and indexed using samtools^48^ and gene expression was quantified as unique read pairs aligned to reference annotated gene features (M82v1.1.1) using HTSeq-count (v0.11.2; parameter --order=pos -- stranded=no --type=exon --idattr=Parent)^85^.

All statistical analyses of gene expression were conducted in R^40^. Differentially expressed genes (DEGs) between the mutants *ssp*, *ssp2*, *ssp ssp2*, and the WT were determined with DESeq2 (v1.34.0)^86^. Raw count data was transformed in DESeqp2 by variant stabilizing transformation (VST). Reproducibility of biological replicates was assessed by hierarchical clustering (method ward.D) and principle component analysis (PCA) using the PCAtools package (v2.6.0) in R^40^. Significantly differentially expressed genes (DEGs) were identified in *ssp* (n=686), *ssp2* (n=180), and *sspssp2* (n=1507) genes with a 1.5-fold change (log_2_FC ≥ 0.58, compared to the WT) and adjusted *p*-value ≤ 0.05 cutoff. Gene normalized z-scores were visualized in heatmaps using pheatmap (v1.10.12) and normalized expression of individual transcripts in transcripts per million (TPM) was plotted using ggplot2.

### Microscopy

Physalis shoot apical meristems manually dissected with a sharp needle and imaged under a Leica stereomicroscope (M205FCA, camera DFC7000T) using the Leica LAS X software (v3.7.0.20979).

### Statistics

Statistical tests and results are reported in Figures and Figure Legends. Significant changes in transactivation assays and in growth phenotypes were determined by one-way ANOVAs followed by post-hoc Tukey’s HSD test at 95% confidence level. Significant differentially expressed genes were determined by Wald’s tests and corrected using the Benjamini and Hochberg (BH) method in DESeq2. Significant differentially bound DAP-seq regions were determined by the Benjamini-Hochberg (BH) method in csaw. Significant changes in normalized read coverage at private DAP-seq regions were determined by performing two-tailed, two-sample t-test. All statistical tests were performed in R.

## DATA AVAILABILITY

The LA1589 genome assembly is available at the Solanaceae Genomics Network (https://solgenomics.net/ftp/genomes/Solanum_pimpinellifolium/LA1589/2020/) and on NCBI under the GenBank accession JBIENU000000000. Raw Nanopore sequence data is available on NCBI SRA under the BioProjects PRJNA1169794, PRJNA607731, and PRJNA557253. Raw Illumina sequence data is available on NCBI SRA under the BioProject PRJNA1069353. The LA1589 SIFT database, deleterious variant predictions, and LA1589 liftoff annotation have been deposited on Zenodo (https://doi.org//10.5281/zenodo.13787654). Sequences of tomato and Arabidopsis bZIP proteins are available on the Plant Transcription Factor Database (PlantTFDB, v5.0, https://planttfdb.gao-lab.org/)^49^. Seeds are available on request from S. Soyk. Data for Supplementary Information is provided as Supplementary Data. Source data are provided with this paper.

## CODE AVAILABILITY

All original code has been deposited on GitHub (https://github.com/soyklab/delvar-2024) and Zenodo (https://doi.org//10.5281/zenodo.13805190)^87^ is publicly available as of the date of publication.

