## Supplementary Figures for "Repairing a deleterious domestication variant in a floral regulator gene of tomato by base editing"

### Supplementary Figure 1

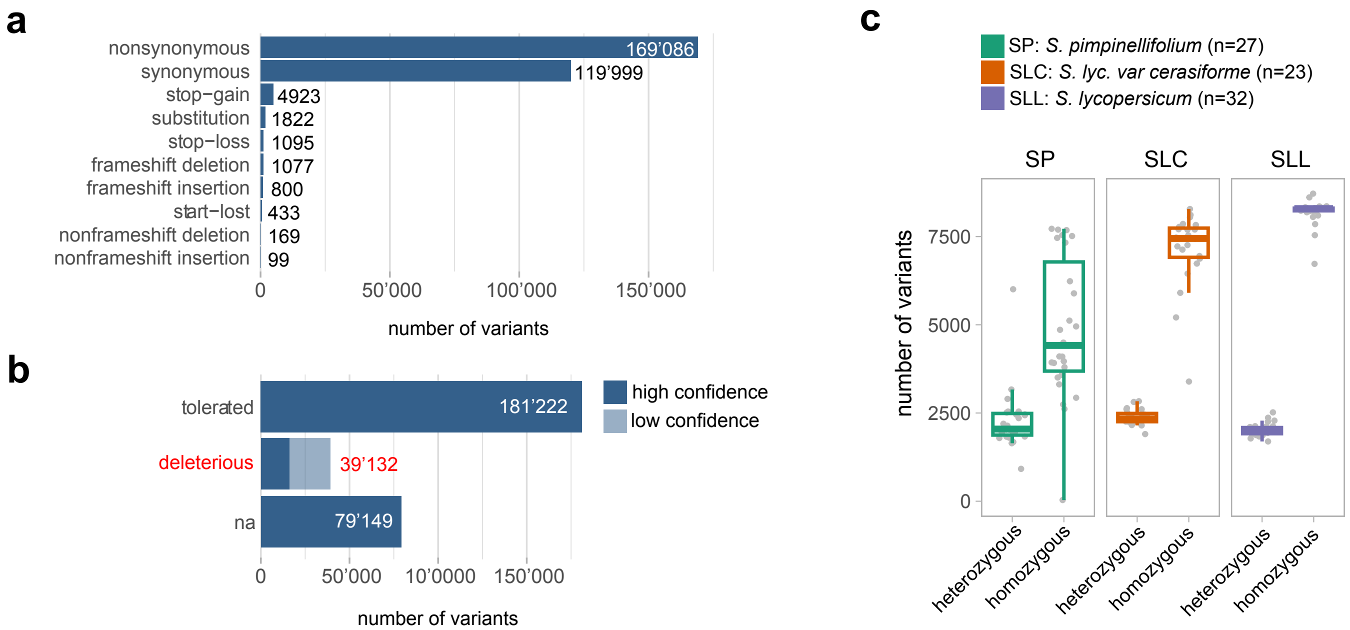


**Supplementary Figure 1: Prediction of deleterious variants in tomato.** **a**, Number of coding sequence variants across a panel of 82 genomes. **b**, Number of non-synonymous variants predicted to be tolerated (sift-score ≥ 0.05), deleterious (sift-score < 0.05), or without prediction (na). Color code indicates confidence of SIFT prediction. **c**, Number of heterozygous and homozygous predicted deleterious mutations in wild (*S. pimpinellifolium*, n=27, in green), landrace (*S. lyc. var. cerasiforme*, n=23, in orange), and domesticated (*S. lycopersicum*, n=32, in purple) tomato genomes. For box plots in (c), the bottom and top of boxes represent the first and third quartile, respectively, the middle line the median and the whiskers the maximum and minimum values.

### Supplementary Figure 2

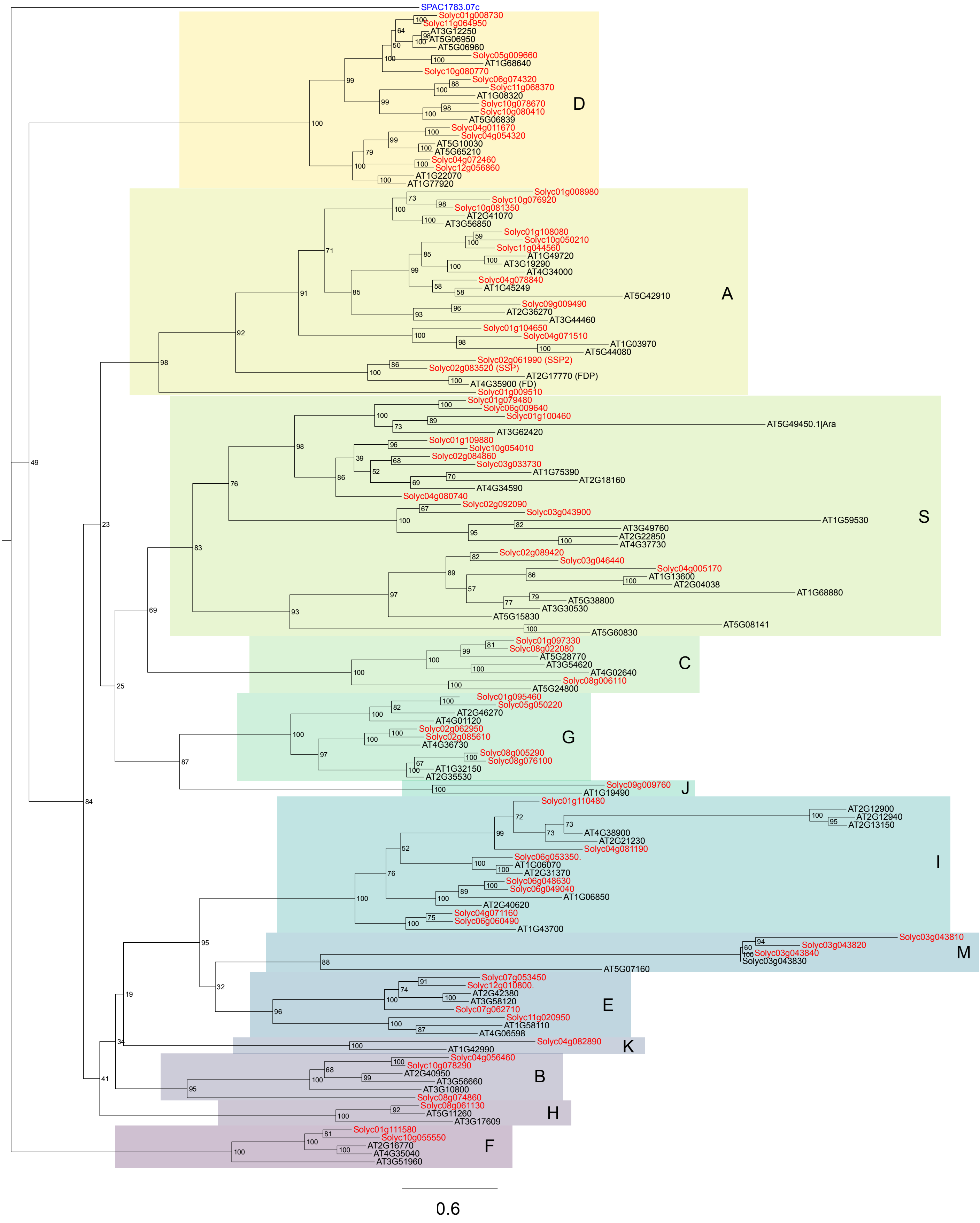


**Supplementary Figure 2: Phylogenetic analysis of the bZIP transcription factor family in Arabidopsis and tomato.** Maximum-likelihood phylogenetic tree constructed with full-length bZIP protein sequences from Arabidopsis (n=74) and tomato (n=70). Arabidopsis and tomato proteins are indicated in black and red font, respectively. The yeast protein Pap1 was used as an outgroup (blue font). Proteins were classified into 13 groups (A-K, M, S) according to the Arabidopsis nomenclature [40]. Numbers represent bootstrap values from 1000 replicates, and scale bar indicates the average number of substitutions per site.

### Supplementary Figure 3

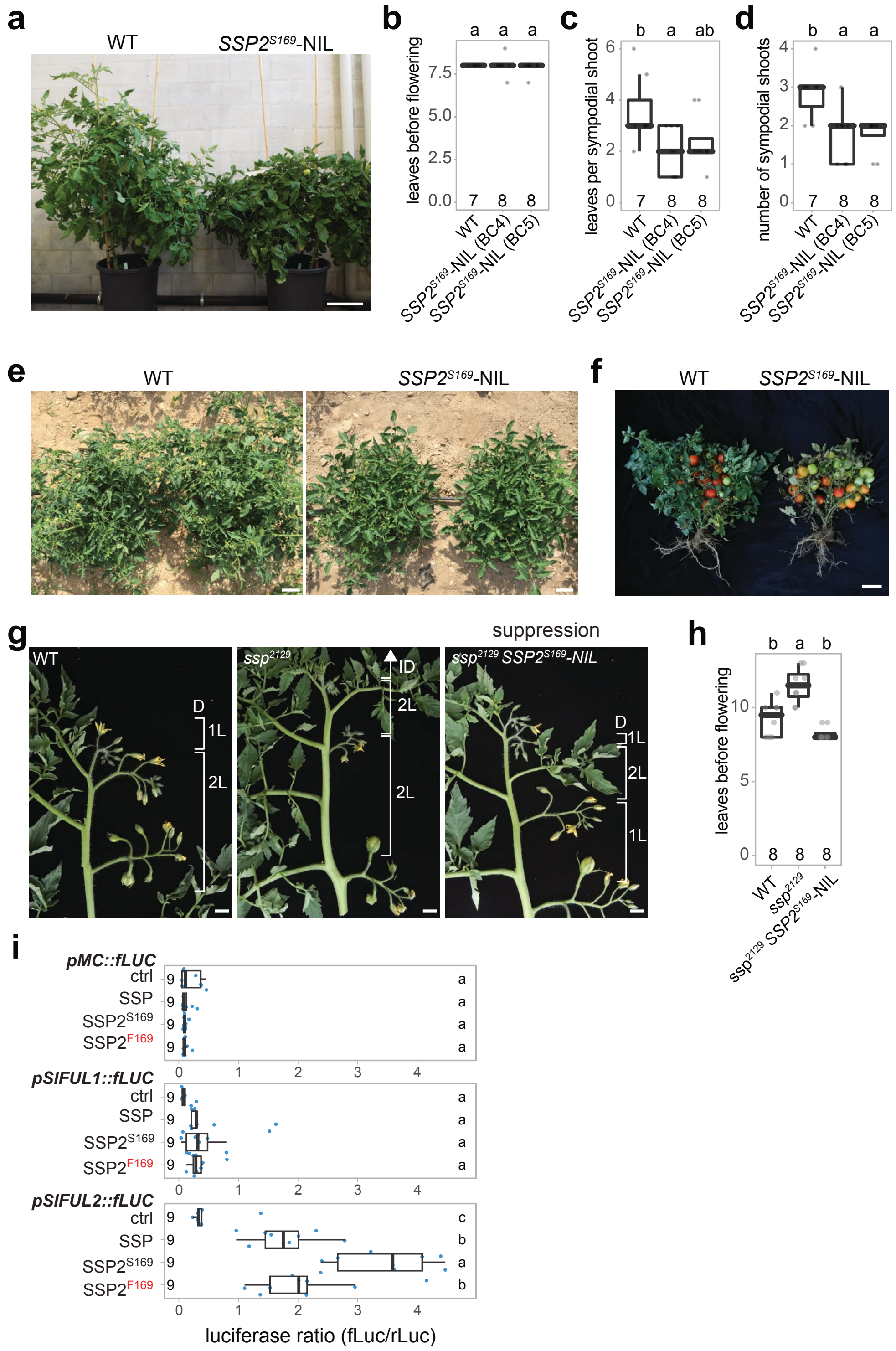


**Supplementary Figure 3: Ancestral *SSP2$^{S169}$* suppresses late flowering and indeterminate growth of *ssp* mutants and shows increased transactivation activity in reporter assays. a**, Representative image of greenhouse-grown wild-type (WT) and *SSP2$^{S169}$*-NIL individual in the determinate M82 background. **b-d**, Quantification of the floral transition (the number of leaves before flowering) on primary (b) and sympodial shoots (c), and the number of sympodial shoot units (d). **e, f,** Representative images of field-grown WT and *SSP2$^{S169}$*-NIL plants at flowering (c) and fruiting (d) stage. **g**, Representative images of detached WT, *ssp$^{2129}$* and *ssp$^{2129}$ SSP2$^{S169}$*-NIL shoots (in the determinate M82 background). D, determinate; ID, indeterminate; L, leaves. **h**, Quantification of the floral transition on the primary shoot for genotypes shown in (e). **i**, Repetition of the reporter assays (**see Fig. 2f**) in tobacco leaves using SSP, SSP2$^{F169}$, and SSP2$^{S169}$ as effectors and firefly Luciferase (fLuc) driven by upstream sequences of *MC* (*pMC::fLUC*), *SlFUL1* (*pSlFUL1::fLUC*), and *SlFUL2* (*pSlFUL1::fLUC*) as reporter. Numbers indicate biological replicates. Ctrl indicates the no effector control (35S::YFP). Numbers at the bottom of the plots of (b) and (f) represent the number of replicate plants and letters at the top of plot (b), (f) and on the right of plot (i) represent the results from pairwise comparisons of means using one-way ANOVA and post hoc Tukey's HSD test with 95% confidence level, respectively. Scale bars indicate 10 cm (a, e, f) and 1 cm (g). For box plots in (c-d, h, i), the bottom and top of boxes represent the first and third quartile, respectively, the middle line the median and the whiskers the maximum and minimum values.

### Supplementary Figure 4

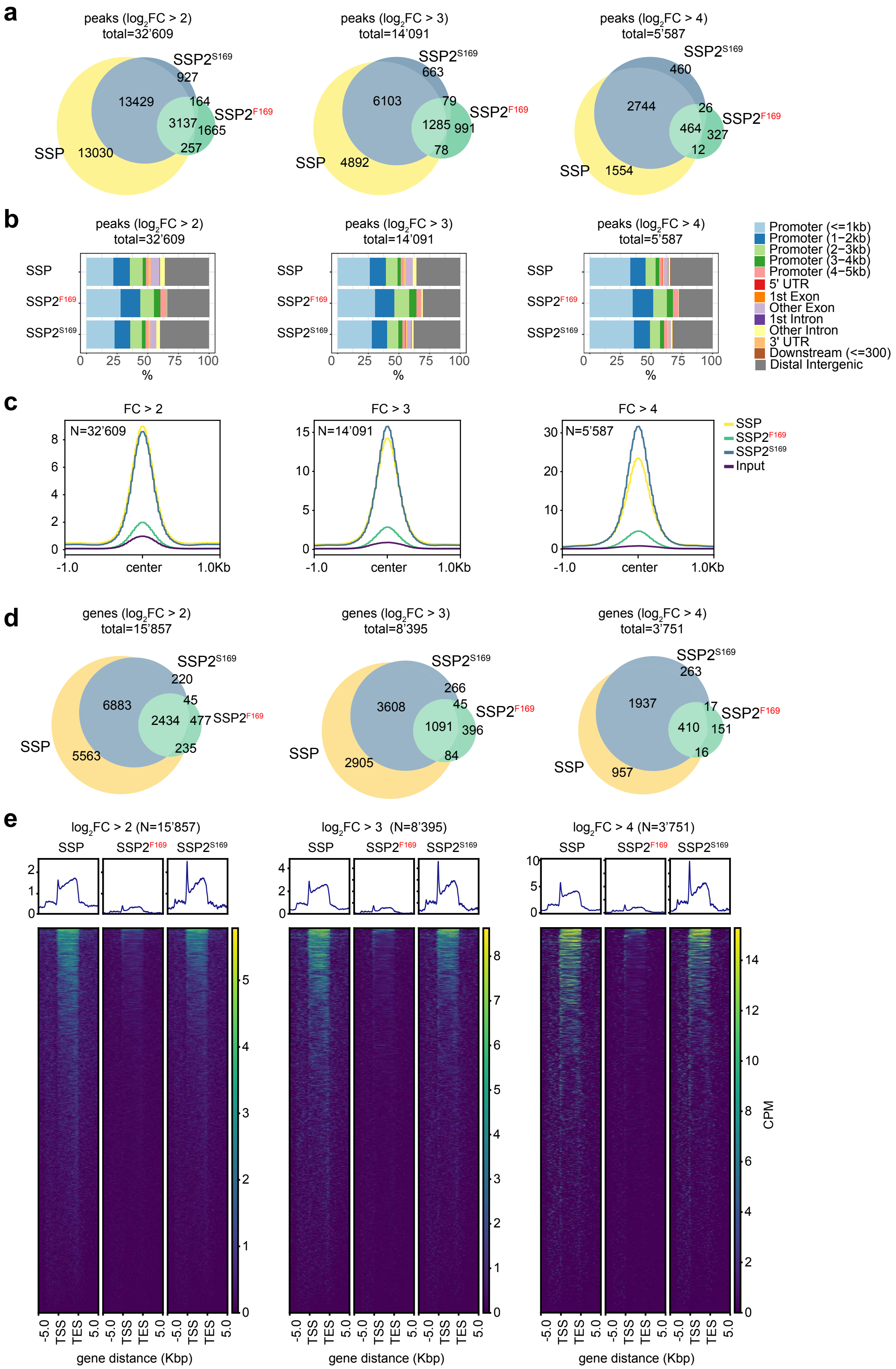

**Supplementary Figure 4: Identification of SSP, SSP2$^{F169}$, and SSP2$^{S169}$ genome-wide binding sites by DAP-seq. a**, Overlap of SSP, SSP2$^{F169}$, and SSP2$^{S169}$ DAP-seq peaks at different significant thresholds ($log_2FC \geq 2, 3, 4$). **b**, Distribution of SSP, SSP2$^{F169}$, and SSP2$^{S169}$ DAP-seq peaks across gene features at different significant thresholds as in (a) **c**, Profiles of normalized read coverage at SSP, SSP2$^{F169}$, and SSP2$^{S169}$ peaks at different significant thresholds as in (a). **d**, Overlap of genes with DAP-seq peaks ≤ 3 Kbp upstream and ≤ 2 Kbp downstream of the transcriptional start site, at different significant thresholds as in (a). **e**, Comparison of SSP, SSP2$^{F169}$, and SSP2$^{S169}$ DAP-seq peaks relative to the transcriptional start (TSS) and end (TES) site of nearby genes, at different significant thresholds as in (a). Top and bottom panels show coverage profiles and heatmaps, respectively.

### Supplementary Figure 5

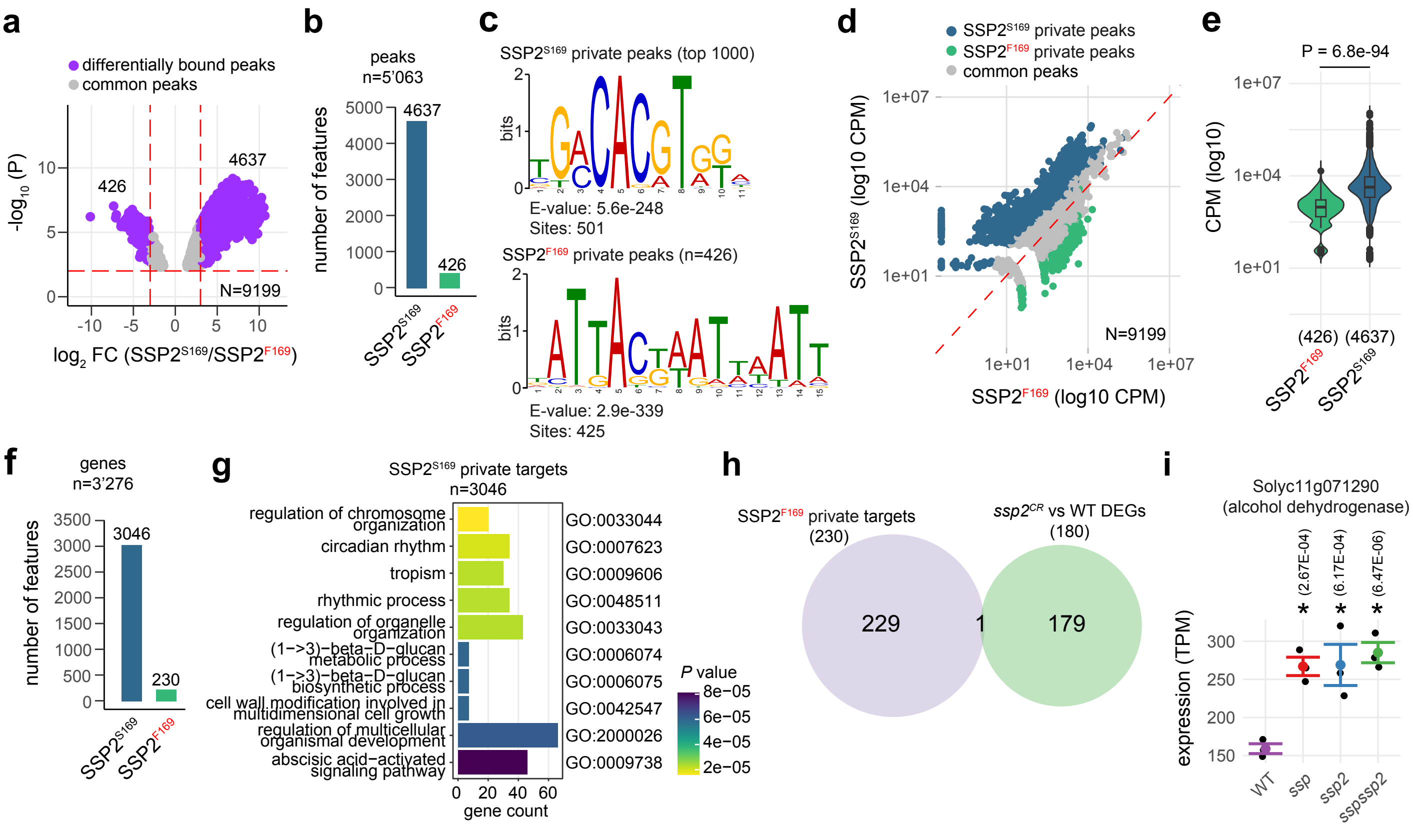


**Supplementary Figure 5: Identification of genomic regions that are differentially bound by SSP2$^{S169}$ and SSP2$^{F169}$. a**, differentially bound (DB) regions (log$_2$FC ≥ 3, FDR ≤ 0.01; by the Benjamini-Hochberg (BH) method in csaw) between SSP2$^{S169}$ and SSP2$^{F169}$ identified by DAP-seq. **b**, number of private DAP-seq regions for SSP2$^{S169}$ and SSP2$^{F169}$. **c**, most-significant motifs identified by *de-novo* motif enrichment analysis of SSP2$^{F169}$ and SSP2$^{S169}$ private DAP-seq regions. **d**, normalized read coverage (CPM, counts per million) at private DAP-seq regions for SSP2$^{S169}$ and SSP2$^{F169}$. **e**, distribution of normalized read coverage at private DAP-seq regions for SSP2$^{S169}$ and SSP2$^{F169}$. *P* represents result from two-tailed, two sample t-test. **f**, number of genes associated with private DAP-seq regions. **g**, the 10 most enriched gene ontology categories for genes associated with private SSP2$^{S169}$ DAP-seq peaks. Note that no enriched categories were detected for SSP2$^{F169}$ private targets. *P* values obtained by Benjamini-Hochberg (BH) method in clusterProfiler. **h**, overlap between SSP2$^{F169}$ private DAP-seq targets and differentially expressed genes (DEGs) in the *ssp2$^{CR}$* mutant compared to the WT identified by RNA-seq. **i**, Normalized expression values (TPM, transcripts per million) for the gene in the overlap in g. Black and colored dots show values from 3 independent, biological replicates and their means, respectively. Error bars indicate standard error. Asterisks (*) indicate significant expression changes (FDR < 0.05) compared to the WT. FDR values (by Wald's tests and corrected using Benjamini and Hochberg (BH) method in DESeq2) are shown. For box plots in (e), the bottom and top of boxes represent the first and third quartile, respectively, the middle line the median and the whiskers the maximum and minimum values.

### Supplementary Figure 6

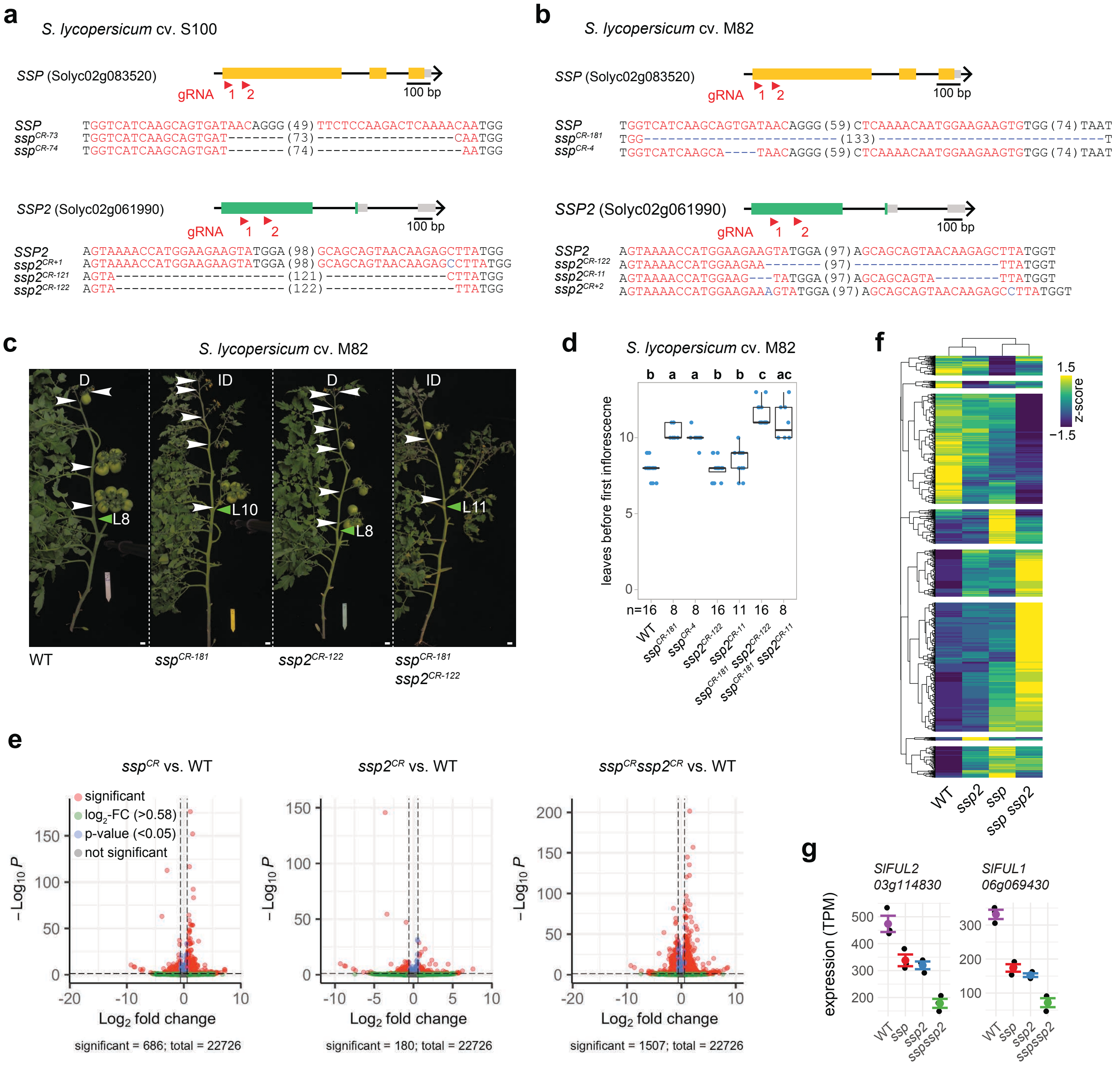


**Supplementary Figure 6: Targeting *SSP* and *SSP2* in two tomato cultivars by CRISPR-Cas9. a,b** CRISPR-Cas9 targeting of *SSP* and *SSP2* in *S. lycopersicum* cv. S100 (a) and cv. M82 (b). Orange boxes, black lines, and grey boxes represent exonic, intronic, and untranslated regions, respectively. Single guide RNAs (sgRNAs) are indicated with red arrowheads. PAM and protospacer sequences are indicated in black and red bold letters, respectively; deletions are indicated with blue dashes; sequence gap length is given in parenthesis. **c**, Representative images WT S100, *ssp*$^{CR}$ and *ssp2*$^{CR}$ single mutants, and *ssp ssp2*$^{CR}$ double mutants. L= leaf number, white arrowheads mark inflorescences. Determinate (D) and indeterminate (ID) shoots are indicated. Scale bars represent 1 cm. **d**, Quantification of the floral transition on the primary shoot for genotypes in (c). N, number of plants. Letters represent the results from pairwise comparisons of means using one-way ANOVA and post hoc Tukey's HSD test with 95% confidence level. **e**, Volcano plots showing differentially expressed genes ($\log_2$ FC > 0.58, FDR < 0.05; by Wald's tests and corrected using Benjamini and Hochberg (BH) method in DESeq2) in *ssp*$^{CR}$ and *ssp2*$^{CR}$ single mutants, and *ssp ssp2*$^{CR}$ double mutants compared to WT (cv. M82). **f**, Heatmap of z-scores showing expression pattern for 1'816 genes that are differentially expressed ($\log_2$ FC > 0.58, FDR < 0.05) in *ssp*$^{CR}$, *ssp2*$^{CR}$ single mutants, and/or *ssp ssp2*$^{CR}$ double mutants in M82. **g**, normalized expression (TPM, transcripts per million) for *SlFUL1* and *SlFUL2* in the WT, *ssp* and *ssp2* single mutants, and the *ssp ssp2* double mutant. Black and colored dots indicate values from 3 independent, biological replicates and their means, respectively. Error bars indicate standard error. For box plots in (d), the bottom and top of boxes represent the first and third quartile, respectively, the middle line the median and the whiskers the maximum and minimum values.

### Supplementary Figure 7

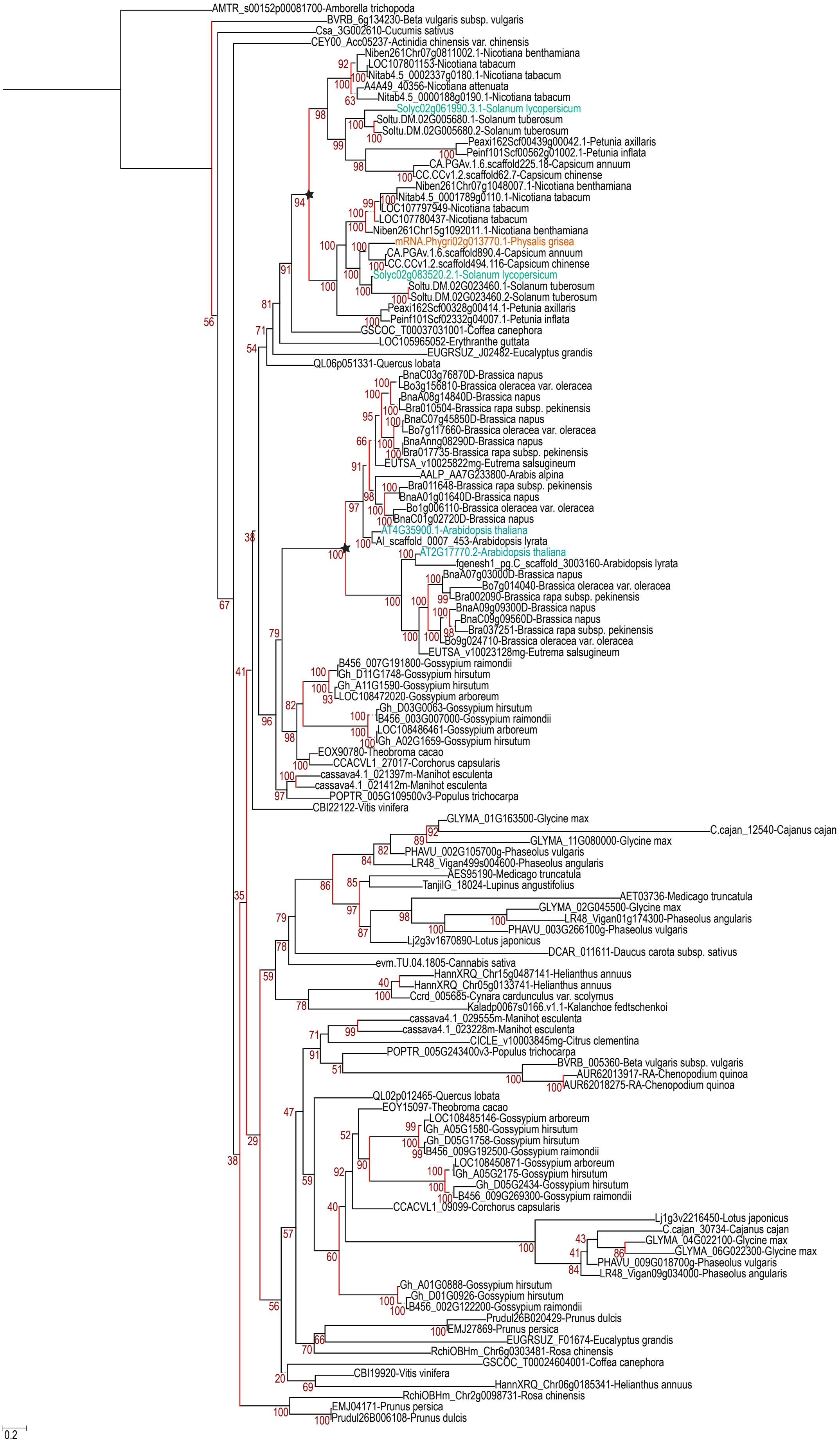

**Supplementary Figure 7: Phylogenetic analysis of SSP homologs in eudicots.** Maximum-likelihood phylogenetic tree constructed with 128 full-length bZIP protein sequences from 51 eudicot species. Tomato, Arabidopsis, and Physalis proteins are highlighted in red, blue, and orange font, respectively. Red branches indicate duplication events, and the two separate duplication events in the *Solanaceae* and *Brassicaceae* are highlighted with stars. Numbers represent bootstrap values from 1000 replicates, and scale bar indicates the average number of substitutions per site.

### Supplementary Figure 8

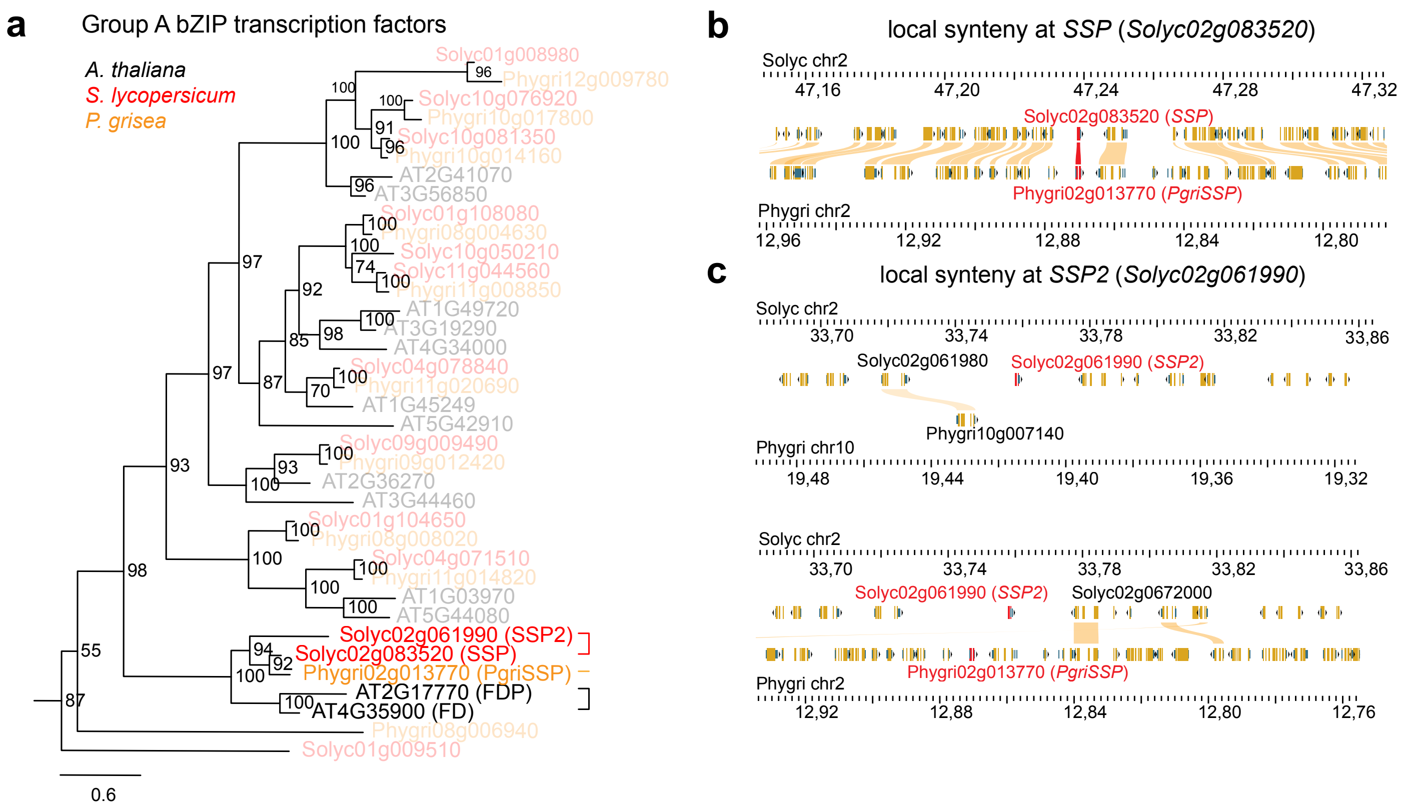


**Supplementary Figure 8: The ortholog of *SSP2* in *Physalis grisea* was lost during evolution. a**. Maximum-likelihood phylogenic tree of the group A bZIP transcription factor family of *A. thaliana*, *S. lycopersicum* and *P. grisea*. Numbers represent bootstrap values from 1000 replicates, and scale bar indicates the average number of substitutions per site. **b,c**, Browser view of synteny analysis of *SSP* (b) and *SSP2* (c) between tomato (cv. S100) and *P. grisea*. Yellow rectangles show annotated genes and yellow streaks link them with their syntenic counterpart. *SSP* and *SSP2* genes are indicated in red. Note the lack of a unique syntenic block for *SSP2* in *P. grisea* in (c).

### Supplementary Figure 9

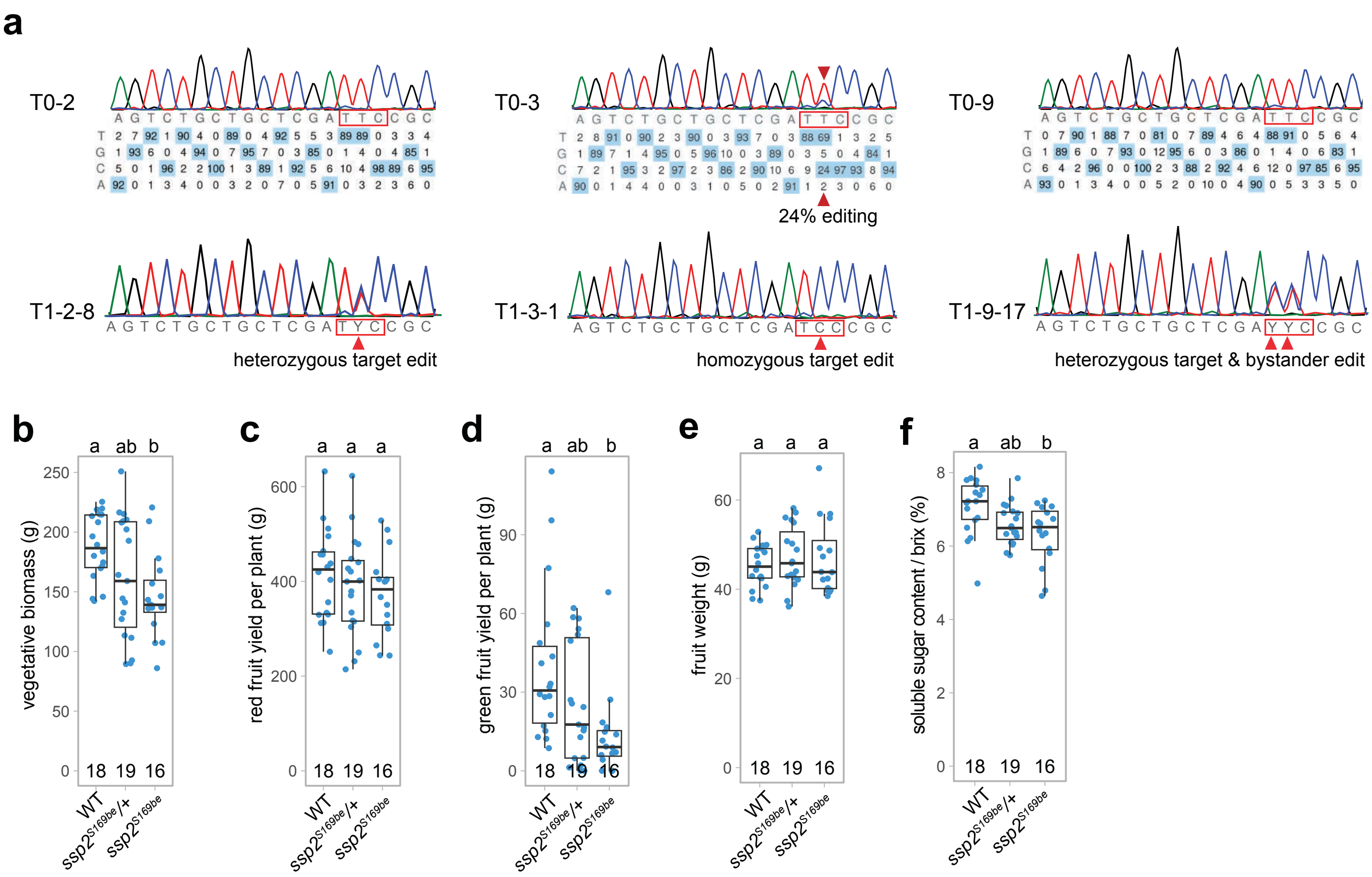


**Supplementary Figure 9: Base-editing of *SSP2* in domesticated tomato and its effect on different tomato yield components. a**, CRISPR base-editing sequencing result of three T0 individuals (upper row) and their T1 progeny (lower row). Note that the target edit was detected in only one T0 individual (T0-3) but in three T1 families. One T1 individual (T1-9-17) was also edited at the bystander adenine. The edited nucleotides are indicated by a red arrowhead. **b-f**, Quantification of the vegetative biomass (b), total red and green fruit harvest (c,d), average fruit weight (e), and average soluble sugar content (brix) (f). The number of plants is indicated in the plots. Letters on top of the plots represent the results from pairwise comparisons of means using one-way ANOVA and post-hoc Tukey's HSD test with 95% confidence level. For box plots in (b-f), the bottom and top of boxes represent the first and third quartile, respectively, the middle line the median and the whiskers the maximum and minimum values.
